# Segmental Acceleration in Protein Condensates

**DOI:** 10.64898/2026.11.24.768996

**Authors:** Keerthivasan Muthukumar, Beata Szała-Mendyk, Dinesh Sundaravadivelu Devarajan, Andrzej Molski, Young C. Kim, Arash Nikoubashman, Jeetain Mittal

**Author notes:** Corresponding authors’. K.M. and B.S.M. contributed equally to this work. Centre of Molecular and Macromolecular Studies Polish Academy of Sciences, Sienkiewicza 112, Lodz, 90-363 Poland (Beata Szała-Mendyk).

## Abstract

Biomolecular condensates are essential to cellular processes and organization, and are typically formed by weak, multivalent interactions among biomolecules, such as intrinsically disordered proteins (IDPs). Because condensates are much more viscous than the surrounding solution, they are generally expected to slow protein motions. While the length-scale-dependent dynamics of IDPs in dilute and dense phases have been studied separately, a systematic comparison of how these dynamics change upon phase separation remains lacking. In this study, we use coarse-grained, explicit-solvent molecular dynamics simulations to compare the segmental dynamics between the dilute and dense phases of the low-complexity domain of the RNA-binding protein Fused in Sarcoma (FUS-LC) and compositionally similar model IDPs of different lengths. Short protein segments, contrary to expectation, relax faster within the condensate than in the dilute phase, whereas segments longer than about one-seventh of the chain relax more slowly. Our results suggest that this crossover arises because chain collapse in the dilute phase slows local relaxation, whereas neighboring chains in the condensate slow chain-scale relaxation. The dilute phase sets both the magnitude of the speed-up and the segment length up to which it extends: dilute-phase relaxation changes with chain length and interaction strength, whereas relaxation within the condensate, at a fixed protein concentration, does not. Slower water dynamics increase the speed-up, whereas removing water eliminates it. Overall, our results suggest that condensates slow global protein motions but accelerate local ones, which may help preserve the liquid-like nature of the condensate’s interior.

## INTRODUCTION

Biomolecular condensates are concentrated assemblies, typically formed by phase separation^1–4^ driven by multivalent interactions^4^ among biomolecules, including intrinsically disordered proteins (IDPs)^5,6^. They are now recognized as central organizers of many cellular functions^7,8^, such as binding and enzymatic modification. These functions may depend on fast IDP motions, which the crowded and viscous condensate interior could slow^9^. IDPs exhibit dynamics spanning multiple length scales, from local backbone fluctuations to segmental and global chain motions^10,11^. Nuclear magnetic resonance (NMR) relaxation studies performed under varying inert and biomolecular crowding conditions demonstrated that IDPs retain fast segmental motions in crowded environments and that both short- and long-range motions depend strongly on the friction of the surrounding solvent^12,13^. Molecular dynamics (MD) simulations showed that the fast normal modes of IDPs within biomolecular condensates exhibit near-identical relaxation across various sequence lengths and degrees of charge segregation^14^. Atomistic simulations of condensates of the low-complexity domain of the RNA-binding protein Fused in Sarcoma (FUS-LC)^15–17^ showed that the proteins remain mobile despite their high concentration^18^, and Martini simulations of FUS-LC and the TIA1 low-complexity domain found that whole chains reconfigure about 20 times more slowly in condensates than in solution^19^. Probe-particle simulations^20^ and single-molecule experiments^21^ further showed that the effective viscosity experienced by small probes and by the chains themselves can be much lower than the bulk viscosity^22,23^.

Considerable effort has been devoted to understanding how IDP sequence features dictate their structure and dynamics in the dilute^24–27^ and dense^14,28–30^ phases *separately*^31,32^. Charge segregation, for instance, modulates these properties differently in each phase; one simulation study showed that increasing charge blockiness results in a coil-to-globule transition and faster chain relaxation in the dilute phase^27^, whereas another study reported the opposite trend within the condensate^28^. Directly comparing the dynamics of IDPs before and after phase separation is essential for understanding how the local protein environment controls the collective properties, functions, and aging of biomolecular condensates^9,21,33–36^. Single-molecule experiments and atomistic MD simulations revealed that, despite the condensate being 300 times more viscous than the dilute phase, chain reconfiguration slows only about threefold upon phase separation, and the number and duration of transient contacts between IDPs remain similar before and after phase separation^21^. Some recent experiments even suggest that local motions along the chain are faster within the condensate than in the dilute phase^37,38^, which is at odds with the expectation that higher viscosity slows all motions. In contrast, static and time-resolved fluorescence anisotropy measurements on other IDPs, including FUS-LC, revealed slower local reorientation in the condensed phase than in the dilute phase^39,40^, and NMR relaxation of the measles virus *N*_TAIL_ domain showed slower motions on all measured timescales^41^. However, these measurements report the reorientation of a fluorescent dye or a backbone bond near a single site, which differs from the relaxation of chain segments of a given length. Despite this progress, it remains unclear how length-scale-dependent IDP dynamics differ between dilute and dense phases.

To address this knowledge gap, we use coarse-grained MD simulations with an explicit solvent to examine how IDP dynamics differ between dilute and dense phases, focusing first on FUS-LC. To generalize our findings, we extend the analysis to simple peptides composed of repeated units of the four-residue motif containing serine (S), tyrosine (Y), glycine (G), and glutamine (Q), with varied total lengths of (SYGQ)*_n_* for *n* = 10, 20, 30, 40, and 50. These simple sequences allow us to avoid complications arising from heterogeneous residue patterns; moreover, S, Y, G, and Q are among the most abundant amino acids in FUS-LC^16,42^, reflecting the high content of polar and aromatic residues characteristic of several phase-separating IDPs, such as hnRNPA1-LC, DDX4-NTD, and TDP-43-CTD^43,44^. Using this minimal model system, we investigate how the motions of chain segments of different lengths change upon condensation. We find that short segments relax faster within the condensate than in the dilute phase, whereas longer segments relax more slowly. Our results suggest that this crossover arises because chain collapse in the dilute phase slows local relaxation, whereas neighboring chains in the condensate slow chain-scale relaxation.

## RESULTS

### Faster Relaxation of Short FUS-LC Segments Within the Condensate than in the Dilute Phase

FUS-LC is the disordered N-terminal region of the RNA-binding protein FUS, rich in serine (SER; S), tyrosine (TYR; Y), glycine (GLY; G), and glutamine (GLN; Q) residues^16,42^ (*Fig. 1A*). FUS-LC consists of *N*_FUS−LC_ = 163 residues and plays a key role in FUS phase separation. To understand how condensation affects IDP dynamics, we simulated FUS-LC in dilute and dense phases at physiological salt concentration using the coarse-grained Martini 3 model with explicit water (*Fig. 1B*; *Methods*). We characterized the dynamics through a normal mode analysis (*Fig. 1C*; *Methods*). We computed the autocorrelation function *c_p_* of normal modes *X_p_* in dilute and dense phases, where the normalized mode number *N*⁄*p* is the characteristic number of residues in the segment described by mode *p* (*Fig. 1C*). As expected, *c_p_* of all modes decayed to zero in both dilute and dense phases at sufficiently long times (*Fig. 1D* and SI Fig. S1), allowing us to estimate the relaxation time *τ_p_* of each mode (*Methods*). We restrict our analysis to segments longer than three residues to probe genuine segmental dynamics rather than local connectivity effects arising from direct bonding or near-neighbor interactions. In both the dilute and dense phases (*Fig. 1E*), shorter segments relax faster than longer ones, consistent with the expectation that longer segments require more monomers to be moved. However, contrary to the general expectation that dense crowded phases exhibit slower overall dynamics, we observed faster relaxation of short segments within the dense phase than in the dilute phase, by more than twofold for the shortest segments (shaded region in *Fig. 1E*). In contrast, segments longer than about 20 residues and the whole chain relax more slowly. To test whether this counter-intuitive behavior is generic to IDPs of similar composition rather than specific to FUS-LC, we next analyzed model peptides of varied length built from a similar residue motif.

**Figure 1:**
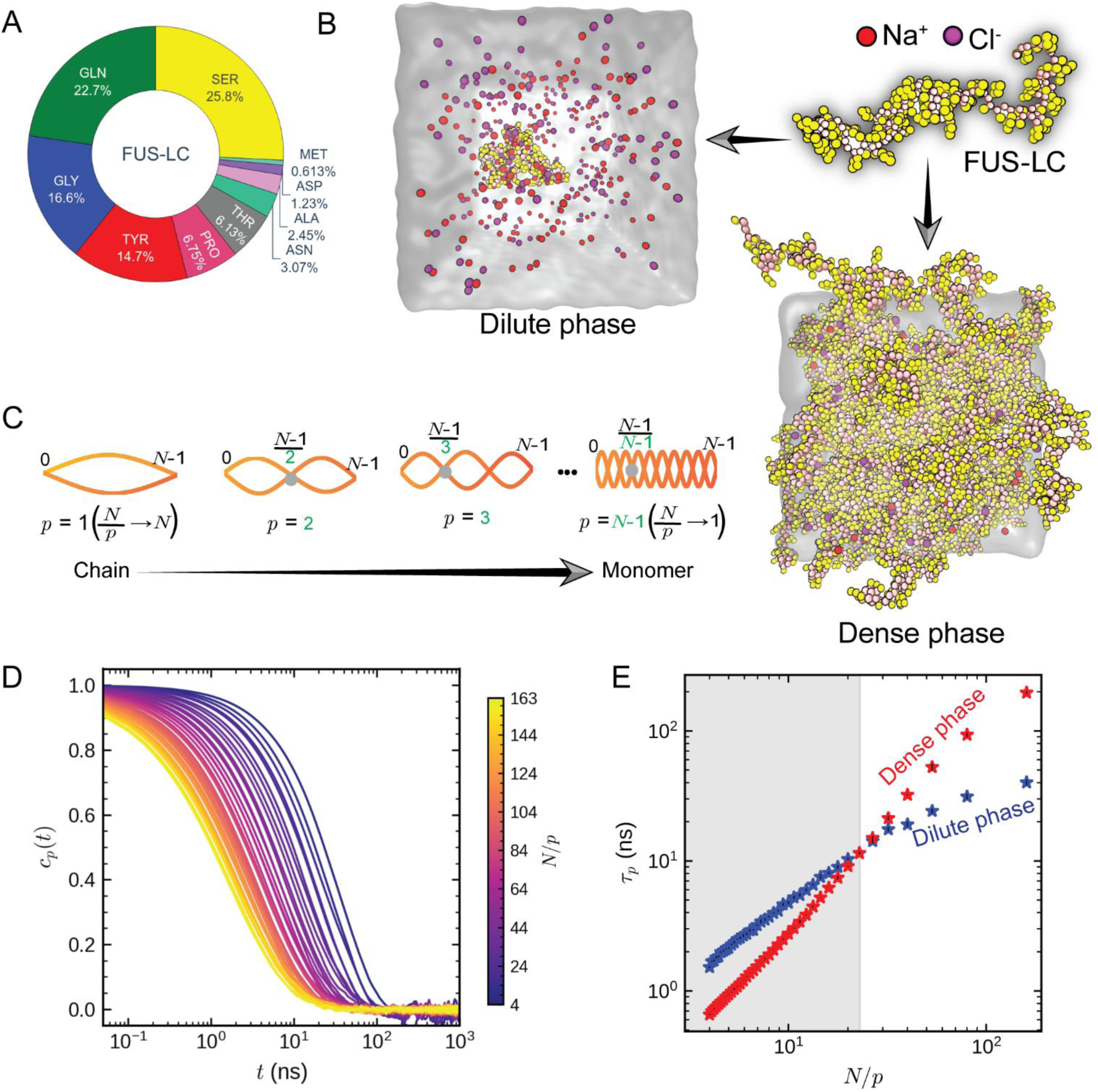
Faster Relaxation of Short FUS-LC Segments Within the Condensate than in the Dilute Phase. (A) Pie chart showing the amino acid composition of the FUS-LC sequence. (B) Simulation snap-shots of the dilute phase (left), a single FUS-LC chain (top right), and the dense phase (bottom right). (C) Schematic representation of normal modes of increasing mode number *p* for a polymer chain of length *N*. Mode *p* = 1 corresponds to whole-chain motion, while higher modes (*p* = 2, 3, . ..) represent increasingly smaller segments down to the monomer scale (*p* = *N* − 1). (D) Normal mode autocorrelation function *c_p_*(*t*) for normalized mode numbers *N*/*p* = 4, 5, … , *N* of the FUS-LC sequence in the dilute phase. Colors, ranging from purple to yellow, represent increasing *N*/*p*. (E) Normal mode relaxation time *τ_p_* of the FUS-LC sequence as a function of normalized mode number *N*/*p* in the dilute and dense phases. The grey shaded region indicates the segments that relax faster within the condensate than in the dilute phase.

### IDP Length Shifts the Dilute-Dense Dynamical Crossover Through Dilute-Phase Relaxation

Given that FUS-LC primarily consists of S, Y, G, and Q amino acids (*Fig. 1A*), we use (SYGQ)*_n_* peptides (*Fig. 2A*) as controllable minimal systems that replicate the salient sequence features of FUS-LC. We first compared the relaxation of FUS-LC with that of an SYGQ peptide of comparable length, (SYGQ)_40_. Despite the reduced sequence complexity of (SYGQ)_40_, its dilute and dense phase relaxations closely follow those of FUS-LC and show the same dilute-dense crossover (*Fig. 2B*). Having demonstrated that the SYGQ peptides can be used as minimal surrogate systems, we next varied the peptide length through the number of SYGQ motifs (*n*) in the sequences and simulated these peptides in dilute and dense phases. Relaxation of all modes in the dilute phase slowed down as the chain length increased, even for segments of the same length (*Fig. 2C*). Power-law fits to the relaxation times (dashed lines in *Fig. 2C*) yielded exponents ranging from 0.94 to 0.98 across different chain lengths, close to the poor-solvent Zimm prediction^45–47^. The Zimm model predicts that the *τ_p_* scales as 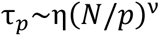, with solution viscosity η and scaling exponent ν (see discussion of segment-level structure below). This relation implies that relaxation times for different *N* in dilute solution should collapse when plotted against *N*/*p*, contrary to our observations. This lack of collapse suggests that the effective dynamical environment experienced by segments depends on the overall chain length. In addition, the curves flatten near the whole-chain relaxation regime. These deviations likely reflect chain collapse in the dilute phase. In longer chains, segments are on average more deeply buried within the compact globule, while long segments relax increasingly through the reorientation of the globule as a whole (*Discussion*). In contrast, in dense phases at a fixed protein concentration of approximately 680 mg/mL (corresponding to the concentration observed in the FUS-LC simulations and approximately 470 mg/mL in real protein mass; *Methods*), the normal mode relaxation times of all (SYGQ)*_n_* peptides collapse onto a single master curve (*Fig. 2D*), consistent with our previous findings for other IDPs consisting of glutamic acid and lysine^14^. The power-law exponent of this master curve is 1.64, between the Zimm value of 1.5 and the Rouse value of 2 for ideal chains^45,48^, as expected when other chains screen hydrodynamic interactions beyond a few residues^46,47,49^. Such screening can be incomplete in dense, water-rich polymer phases^50,51^. Because the dense-phase curve is steeper than the dilute-phase curves, the two cross at an intermediate segment length. Analysis of the ratio 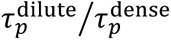 reveals that increasing *N* not only increases the length of the segments that relax faster within the condensate ((*N*⁄*p*)_crossover_), but also increases the 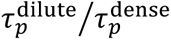 ratio for any given *N*⁄*p* (*Fig. 2E*). This crossover length grows roughly in proportion to chain length, so the crossover always falls near the sixth to eighth normal mode, about one-seventh of the chain (*Fig. 2E*, inset). This effect arises mainly from the variation in dilute phase relaxation (*Fig. 2C*), as the dense phase relaxation is unchanged (*Fig. 2D*). Taken together, these results show that increasing the IDP length systematically shifts the dilute-dense dynamical crossover towards longer segments, while also amplifying the relative acceleration of short-segment relaxation within the condensate. Since the explored range of *N* covers most naturally occurring IDPs^5,52^, this length dependence should apply to many IDPs of similar composition. Because protein conformations differ in dilute and dense phases^53,54^, we next asked whether the crossover in dynamics has a structural counterpart at segmental and global scales.

**Figure 2:**
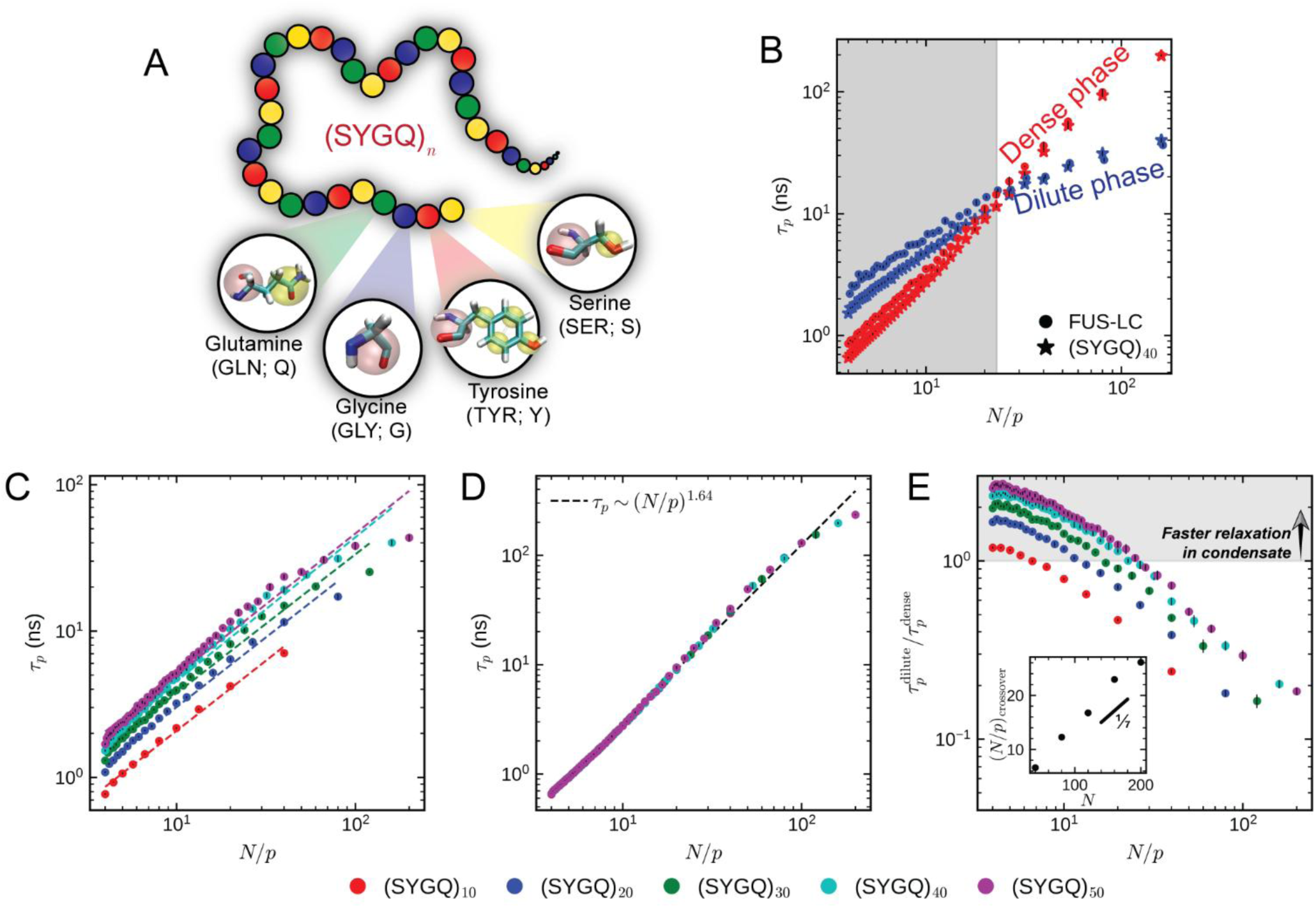
IDP Length Systematically Shifts the Dilute-Dense Dynamical Crossover Toward Longer Segments by Slowing Dilute-Phase Relaxation. (A) Schematic representation of a (SYGQ)_n_ peptide. Serine, tyrosine, glycine, and glutamine residue representation in both the all-atom and coarse-grained Martini model is shown. (B) Normal mode relaxation time *τ_p_* of FUS-LC and (SYGQ)_40_ sequences as a function of normalized mode number *N*/*p* in the dilute and dense phases. *τ_p_* of (SYGQ)_n_ sequences as a function of normalized mode number *N*/*p* in the (C) dilute phase and (D) dense phase. Dashed lines show power-law fits. (E) Ratio 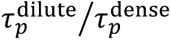 of dilute and dense phase *τ*_p_ as a function of *N*/*p* for (SYGQ)_n_ sequences. The grey shaded region in (E) indicates the segments that relax faster within the condensate than in the dilute phase. The inset in (E) shows the crossover segment length as a function of chain length.

### Segmental Structure Does Not Mirror the Dilute-Dense Dynamical Crossover

We characterize the conformation of the IDPs through their radius of gyration 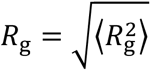 in dilute and dense phases. Irrespective of the SYGQ sequence length, *R*_g_ increases from the dilute to the dense phase, consistent with previous studies^54–56^ (*Fig. 3A*). *R*_g_ scales with *N* as *R*_g_∼*N^v^*, with *ν* = 0.34 in the dilute phase and 0.53 in the dense phase (dashed lines in *Fig. 3A*), consistent with previous studies^14,54^. These exponents indicate that the SYGQ peptides are collapsed in the dilute phase (poor solvent), while they exhibit nearly ideal-chain scaling in the dense phase, likely due to the screening of nonbonded interactions. This interpretation is further supported by the relationship between the root-mean-square end-to-end distance *R*_e_ and *R*_g_, with 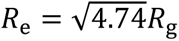 in the dilute phase and 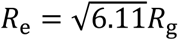 in the dense phase (*SI Fig. S2*). The latter closely agrees with the Gaussian-chain prediction 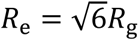. Having established that the chains expand upon phase separation, we next examined whether the crossover seen for the segmental dynamics is mirrored by a corresponding crossover in the segmental structure.

**Figure 3:**
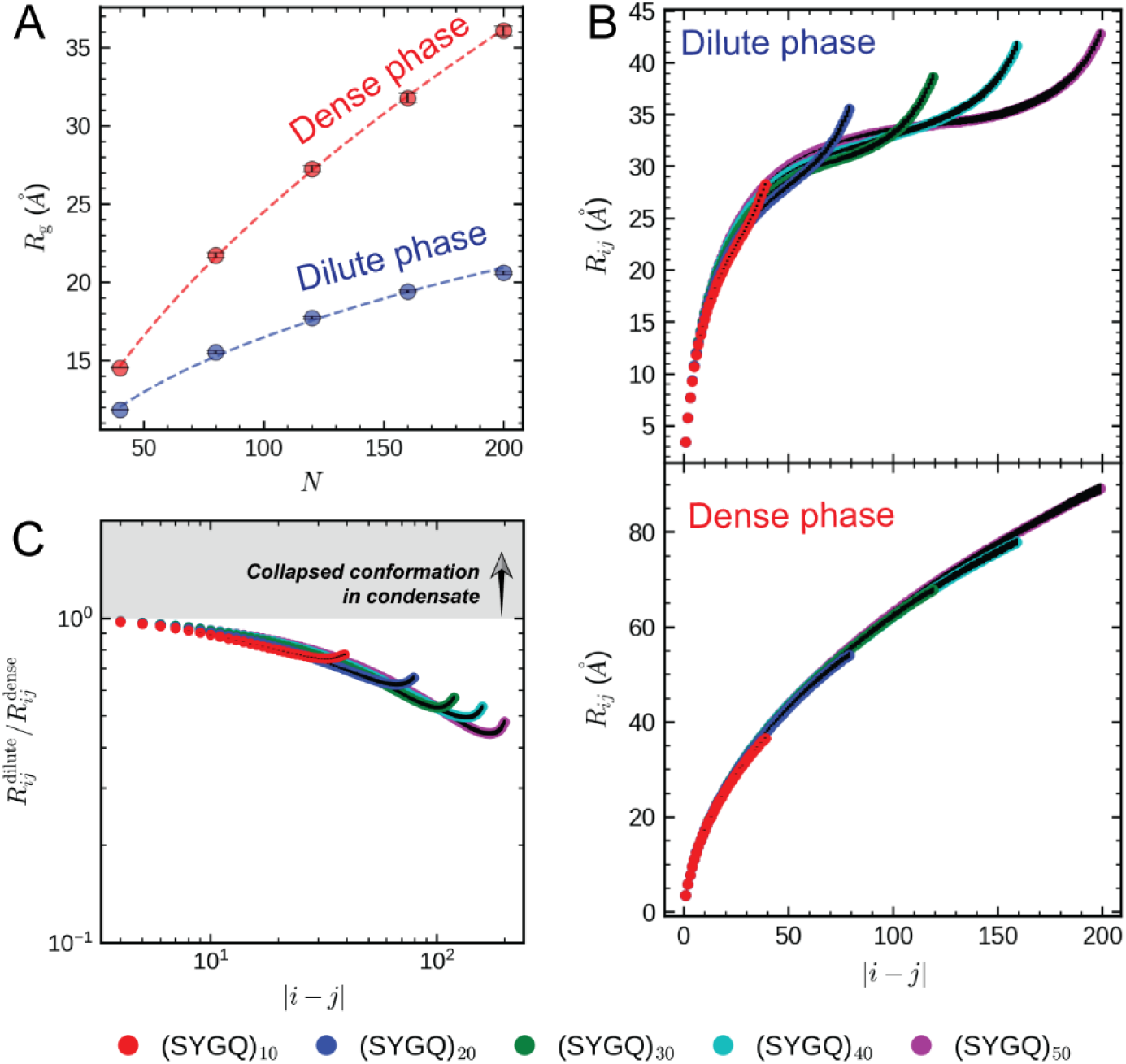
Phase Separation Leads to Large-Scale Chain Expansion While Largely Preserving Short-Segment Structure. (A) Radius of gyration *R_g_* of SYGQ peptides as a function of chain length *N* in dilute and dense phases. Dashed lines show power-law fits. (B) Interresidue distance *R_ij_* versus separation distances |*i* − *j*| for SYGQ sequences in dilute (top) and dense (bottom) phases as a function of *N*. (C) Ratio 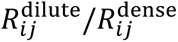 of dilute and dense phase *R_ij_* as a function of sequence separation for (SYGQ)_n_ sequences. The grey shaded region in (C) marks segments that would be more compact in the condensate than in the dilute phase.

To this end, we characterize the segmental conformation via the root-mean-square interresidue distance 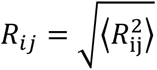 in dilute and dense phases. In both phases, *R_ij_* increases with increasing segment length |*i* − *j*| for all sequences (*Fig. 3B*). In the dilute phase, the interresidue distance stays near 30-35 Å for widely separated residues, as expected for a collapsed chain, whose size limits the distance between any two residues. The sharp rise at the largest separations is likely a chain-end effect. Notably, the curves in the dense phase collapse onto a single master curve, suggesting minimal chain-end effects and a high degree of self-similarity for these (SYGQ)*_n_* sequences. Consistent with near-theta conditions, the end-to-end distance distributions in the dense phase are well described by the empirical Mazur form^14,57^ 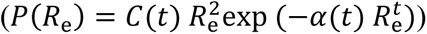 with fitted exponents *t* that approach the Gaussian limit of *t* = 2 for longer chains; deviations from this limit are primarily observed for the shorter chains, where finite-size effects are expected (*SI Fig. S3*). Next, to check whether there is a crossover in the segmental structure, we calculated the ratio 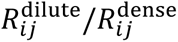, which never exceeded one (*Fig. 3C*), indicating that all segments expand upon phase separation, although the shortest segments expand only slightly (ratio close to one), as expected for segments shorter than a thermal blob, which are nearly ideal in any solvent^46^.

Together, these results show that phase separation causes chain expansion across all length scales. Because short segments have nearly the same size in both phases, their faster relaxation in the condensate is not caused by a change in their structure. Notably, while the overall chain size grows strongly with increasing SYGQ sequence length, the dense phase segmental structure and normal mode relaxation remain invariant, indicating that local dynamics within the condensate do not depend on IDP length at a fixed protein concentration. This behavior raises the question of whether changing the protein-protein and protein-water interactions, which set how collapsed the chains are in the dilute phase, changes the speed-up.

### Interaction Strength Sets the Speed-Up Through Dilute-Phase Chain Collapse

The two main components of the biomolecular condensate environment are the neighboring proteins and water^58,59^. To test how interactions with these two components affect the speed-up within the condensate relative to the dilute phase, we tuned the protein-protein (PP) and protein-water (PW) interactions^60^ at a fixed condensate concentration (corresponding to the concentrations observed in the FUS-LC simulations; *Methods*). In the following, larger *λ_PP_* and *λ_PW_* values correspond to stronger protein-protein and protein-water interactions, respectively, and the interaction-strength variations considered here are small perturbations (< 5%) around the original Martini interaction parameters. Decreasing the protein-protein interactions or increasing the protein-water interactions weakens the net attraction between protein segments. As a result, the chain expanded in the dilute phase (*Fig. 4A* and *4D; SI Fig. S4A* and *S4B*). This structural change is accompanied by a faster relaxation of all normal modes in the dilute phase (*Fig. 4B* and *4E*). Analogous changes in *λ_PP_* and *λ_PW_* in the dense phase resulted in a much smaller chain and segmental expansion (Insets in *Fig. 4A* and *4D; SI Fig. S5A* and *S5B*), consistent with the screening of nonbonded interactions in semidilute polymer solutions^46,47^. However, at this fixed concentration, the relaxation of the normal modes within the dense phase remained unaffected, with data from all curves collapsing onto a single master curve (Insets in *Fig. 4B* and *4E*). This pronounced variation in the dilute phase, with negligible variation in the dense phase, lowered the 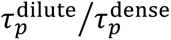 ratio relative to the original Martini parameters (*λ_PP_* = *λ_PW_* = 1) as *λ_PP_* is decreased or *λ_PW_* is increased (*Fig. 4C* and *4F*). For the strongest protein-water interaction tested, for example, the speed-up of the shortest segments falls from about 2.3-fold to about 1.2-fold, and the crossover moves from about 22 to about 7 residues (*Fig. 4F*). At this interaction strength, the dilute chain is nearly as expanded as in the condensate (*R*_g_ about 31 vs 35 Å; *SI Figs. S4B* and *S5B*), yet long segments still relax about three times more slowly in the condensate (*Fig. 4F*). Collapse thus accounts for most of the short-segment speed-up, whereas the slower relaxation of long segments is a property of the condensate.

Thus, changing the PP and PW interactions alters chain and segmental conformations in both phases, but, at a fixed protein concentration, changes only the dilute-phase relaxation. These findings show that the magnitude and extent of the speed-up are set by how collapsed the chain is in the dilute phase. Because the same net attraction drives chain collapse and phase separation^24,61^, we expect the speed-up to grow with the driving force for phase separation at a similar condensate concentration, and its exact magnitude depends on the interaction parameters of the model. As biomolecular condensates retain a substantial water fraction, we next examined whether solvent dynamics, particularly the dynamics of water beads, affect the dynamical crossover between dilute and dense phases.

**Figure 4:**
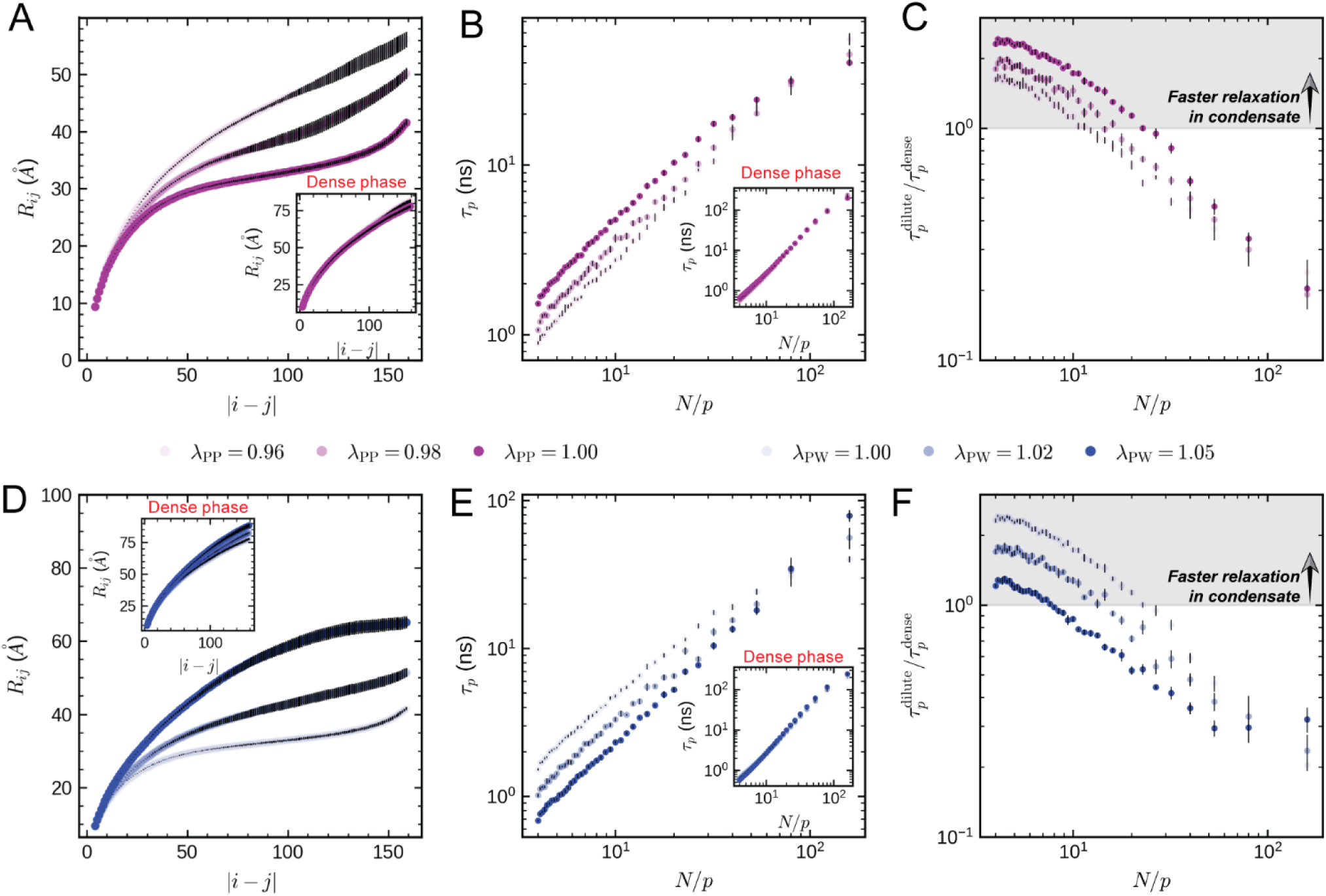
Interaction-Driven Expansion of the Dilute-Phase Chain Shrinks the Speed-Up. (A) Interresidue distance *R_ij_* versus separation distances |*i* − *j*| of (SYGQ)_40_ sequence in dilute and dense (inset) phases for different protein-protein interaction strength *λ_PP_* values. (B) Normal mode relaxation time *τ_p_* of (SYGQ)_40_ sequence as a function of normalized mode number *N*/*p* in the dilute and dense (inset) phases for different *λ_PP_* values. (C) Ratio 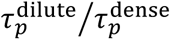 of dilute and dense phase *τ*_p_ as a function of *N*/*p* for (SYGQ)_40_ sequence for different *λ_PP_* values. The grey shaded region indicates the segments that relax faster within the condensate than in the dilute phase. Similar data for different protein-water interaction strength *λ_PW_* values are shown in panels (D)-(F). Dense-phase simulations are at a fixed protein concentration of about 680 mg/mL.

### Dilute-Phase Relaxation Is More Sensitive to Water Dynamics

Similar to the cell itself, water constitutes a major portion of biomolecular condensates^58,59^ and shapes IDP dynamics both in solution and within condensates^62,63^. In our dense phase simulations, water accounts for roughly 60 vol% of the condensate environment. To understand the role of water in the faster segmental motion within the condensate, we systematically varied the water mass in the (SYGQ)_40_ system, in both the dilute and dense phases (*Fig. 5A*; *Methods*). Scaling water mass offers a way to change the timescale of the solvent motion (and hence the viscosity, which is expected to scale as the square root of the water mass), without affecting the static system properties^64–68^. We found that increasing the mass of the water particles resulted in a systematic slowdown of all normal modes in both the dilute and dense phases (*Fig. 5B*). However, this effect is more pronounced in the dilute phase, likely due to its much higher water fraction, so that the 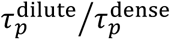 ratio increased with increasing water mass for all modes (*Fig. 5C*). Consequently, the crossover segment length also increased (inset in *Fig. 5C*). Together, these results show that solvent dynamics strongly modulate the relative segmental relaxation between dilute and dense phases.

To probe the coupling between segmental motion and solvent dynamics, we examined how the relaxation time of each normal mode scales with water mass. We fitted the relaxation time of each mode to a power law in water mass, 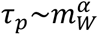, where *α* characterizes the sensitivity of the dynamics to solvent motion, with an exponent of 0.5 expected if relaxation follows the solvent viscosity, and smaller values indicating a weaker dependence, often attributed to internal friction^65,69^. In both phases, the proteins are coupled to water, with exponents about 0.32-0.36 in the dense phase and about 0.36-0.5 in the dilute phase (*Fig. 5D*). Thus, the dense-phase segmental relaxation is less sensitive to water motion, with little change across segment lengths, while the relaxation of long segments in the dilute phase follows the solvent viscosity almost exactly. Therefore, increasing the water mass results in a stronger slowdown in the dilute phase compared to the dense phase, explaining why the ratio 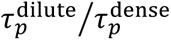 increases with water mass (*Fig. 5C*). Notably, while the relaxation times themselves exhibit a dilute-dense crossover, the exponent does not. Because the condensate has the lower exponent, internal friction makes up a larger share of its friction, yet its short segments relax faster; internal friction therefore does not explain the speed-up.

**Figure 5:**
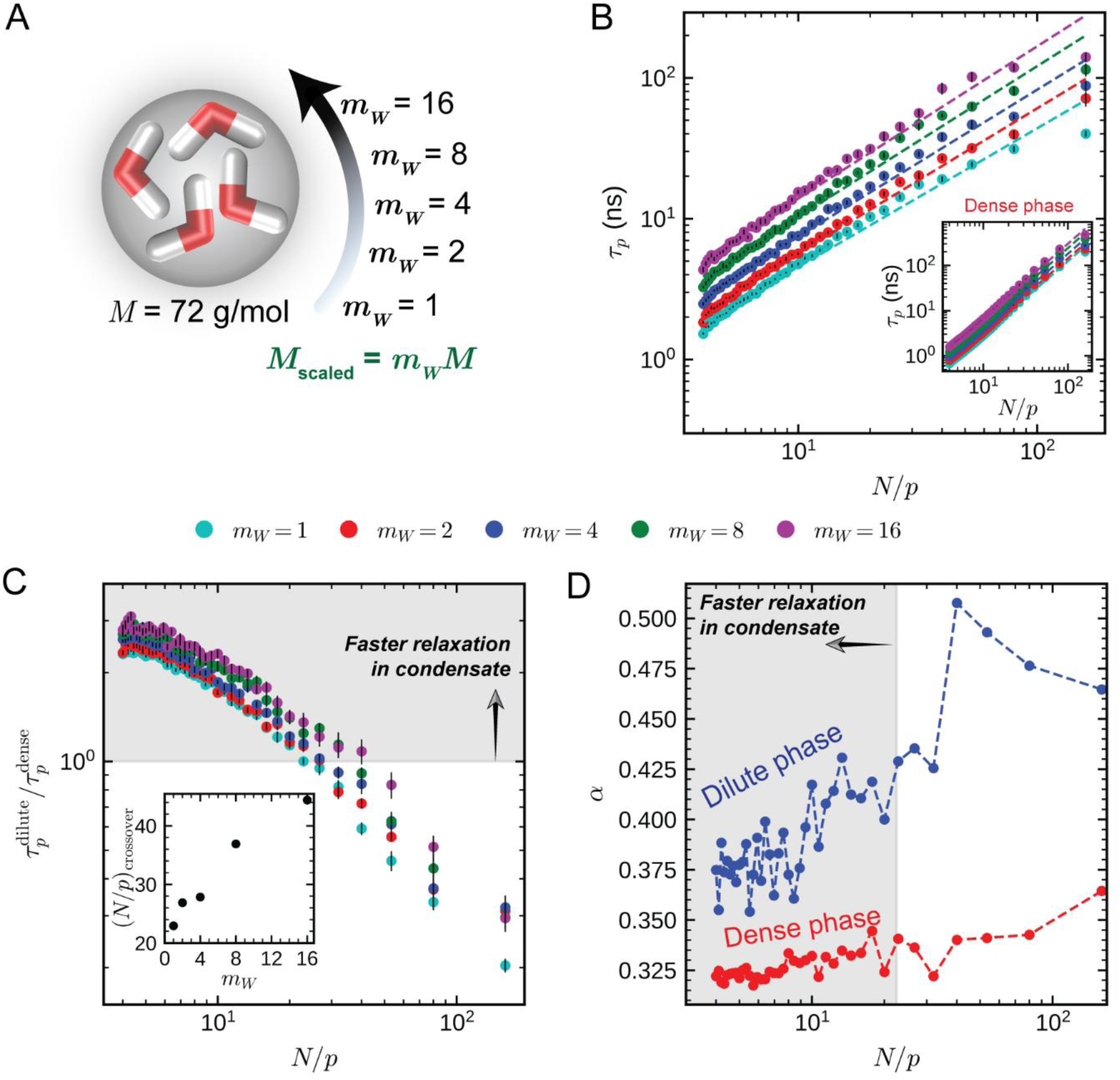
Dilute-Phase Relaxation Is More Sensitive to Water Dynamics. (A) Schematic representation of a standard water bead in the Martini model, where each coarse-grained Martini water has a mass of 72 g/mol and corresponds to 4 real water molecules. Water mass is scaled to a value of *M_scaled_* by multiplying the actual mass by a factor *m_W_* in the simulation. (B) Normal mode relaxation time *τ_p_* of (SYGQ)_40_ sequence as a function of normalized mode number *N*/*p* in the dilute and dense (inset) phases for different *m_W_* values. (C) Ratio 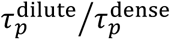 of dilute and dense phase *τ_p_* as a function of *N*/*p* for (SYGQ)_40_ sequence for different *m_W_* values. The inset shows the crossover segment length as a function of *m_W_*. (D) Power law exponent *α* of *τ_p_* scaling with *m_W_* as a function of *N*/*p*. An exponent of 0.5 corresponds to relaxation that follows the solvent viscosity. The grey shaded region in (C) and (D) indicates the segments that relax faster within the condensate than in the dilute phase.

### Absence of Water Eliminates the Dilute-Dense Segmental Dynamics Crossover

Given the strong role of water dynamics in the dilute-dense dynamical crossover, we next asked whether water is required at all. To answer this question, we removed all water particles while retaining ions in both the dilute and dense phases of the (SYGQ)_40_ system. Eliminating the explicit solvent reversed the trend entirely: relaxation became considerably faster in the dilute phase and slower in the dense phase (*Fig. 6A*), driving the dilute-to-dense ratio 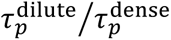 far below one across all modes (*Fig. 6B*). Notably, this acceleration of dilutephase dynamics in the water-free systems occurs despite the chain adopting a more collapsed conformation (*SI Fig. S6A* and *S6B*), which would be expected to increase intrachain friction. In the absence of explicit water, the relaxation dynamics are governed by thermostat coupling (*Methods*) and the remaining nonbonded interactions, including those between proteins and ions. To assess the contribution of the latter, we performed additional simulations in which ions were also removed, leaving only the protein chains (*Methods*). While removing ions decreased the segmental relaxation times in both phases (*SI Fig. S7A* and *S7B*), it further reduced the dilute-to-dense ratio 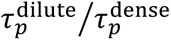, particularly for shorter segments (*SI Fig. S7C*). The ratio remained below one across all segment lengths, indicating that the loss of faster segmental relaxation within condensates upon water removal does not depend on the presence of ions.

Because removing water strongly accelerates dilute phase relaxation, one might similarly expect the water-free condensate to also relax faster if water acted mainly as a viscous drag in both phases. However, the smaller water-mass scaling exponent already showed that dense phase relaxation is less sensitive to solvent motion (*Fig. 5D*). Consistent with this weaker solvent dependence, removing water does not accelerate condensate relaxation and instead leads to considerably slower dynamics in the dense phase. Because short segments in the water-free condensate relax more slowly than even those in the hydrated dilute phase (*Fig. 6A*), the loss of the speed-up is unlikely to depend on the thermostat setting and more likely reflects the stronger influence of nonbonded interactions between protein chains.

**Figure 6:**
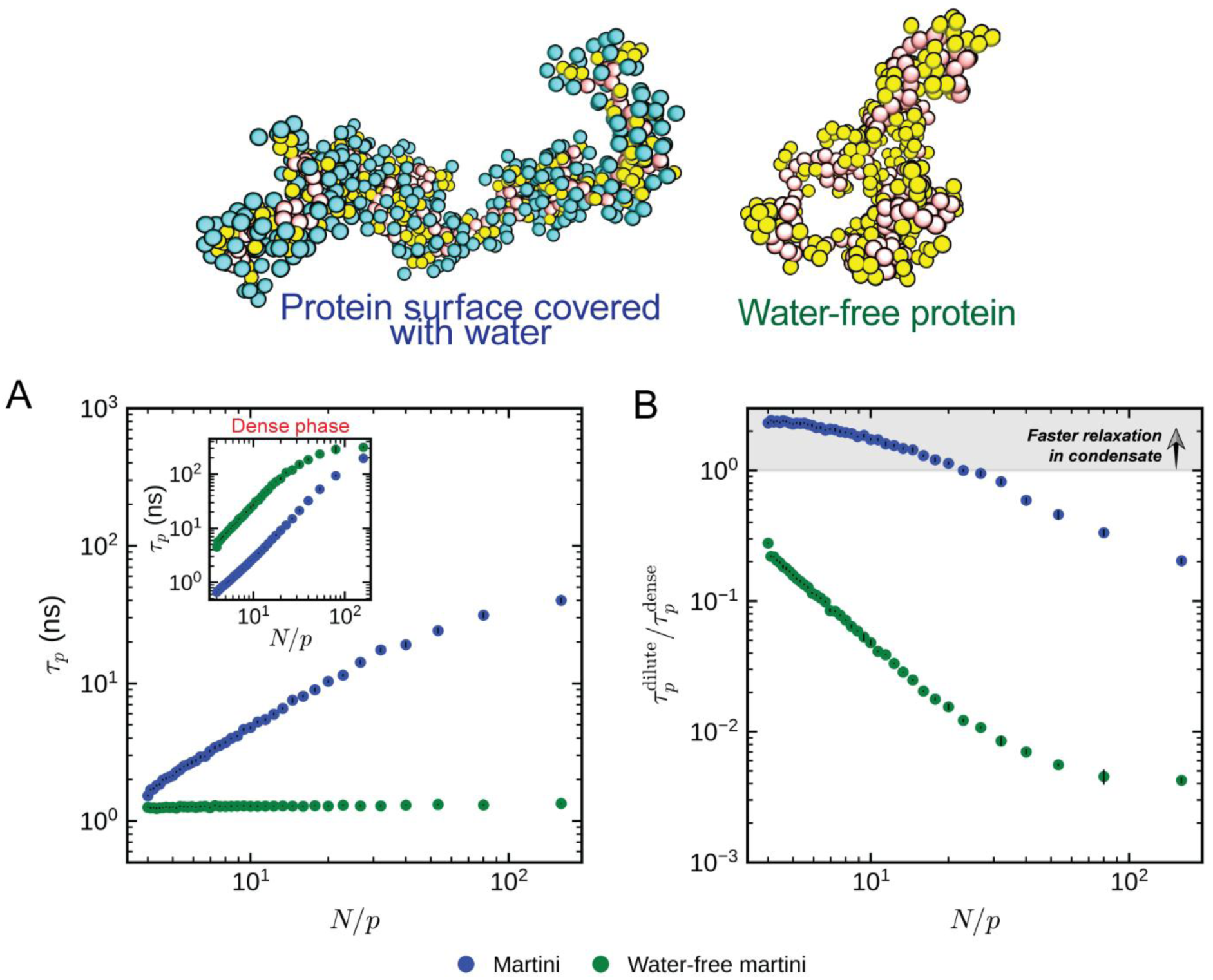
Water-Free Martini Simulations Show That Water Is Required for Faster Segmental Relaxation Within the Condensate. (A) Normal mode relaxation time *τ_p_* of the (SYGQ)_40_ sequence as a function of normalized mode number *N*/*p* in the dilute and dense (inset) phases for hydrated and water-free Martini simulations. (B) Ratio 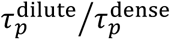of dilute and dense phase *τ_p_* as a function of *N*/*p* for (SYGQ)_40_ sequence in hydrated and water-free Martini simulations. The grey shaded region in (B) indicates the segments that relax faster within the condensate than in the dilute phase.

Moreover, the removal of water does not manifest itself in the same way across all modes: in the dilute phase, the difference between the relaxation times in the solvated and solvent-free simulations increases with increasing *N*/*p* (i.e., for longer segments), whereas the gap decreases in the dense phase (*Fig. 6A*). This non-trivial, mode-dependent rescaling underscores that the degrees of freedom of the solvent and protein cannot be separated by a single global time-scaling factor, and that caution is warranted when mapping coarse-grained simulation timescales to experimental observables^70–72^.

To rule out the possibility that this reversal arises from retaining the volume and protein concentration of the hydrated system after water removal, we additionally re-equilibrated the solvent-free dense phase, allowing its density and chain conformation to readjust. Upon re-equilibration, the solvent-free dense phase contracted, resulting in a concentration increase from ∼680 to ∼1596 mg/mL. At the same time, the chains became more compact (*SI Fig. S8A* and *S8B*), yet the 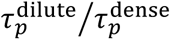 ratio remained below one at all segment lengths examined (*SI Fig. S8C* and *S8D*). Thus, the loss of the crossover upon water removal persists despite the increase in protein density, indicating that it is not an artifact of retaining the initial simulation volume. Together, these results show that explicit water particles are required for the faster segmental relaxation within condensates: without water, the condensate becomes a dry, densely packed protein phase in which all segments relax more slowly than in the dilute phase.

## DISCUSSION

Biomolecular condensates are typically viewed as crowded and highly viscous environments that uniformly slow molecular motion. However, our results show that this picture breaks down at sufficiently small length scales, reminiscent of the probe-size-dependent effective viscosity observed in polymer solutions and condensates^20–23^. Using coarse-grained molecular dynamics simulations of FUS-LC and simplified SYGQ polypeptides, we find that short protein segments relax faster inside the condensate than in the dilute phase, whereas longer segments relax more slowly, showing that phase separation reorganizes protein dynamics in a strongly length-scale-dependent manner. Our findings could help to better interpret experimental measurements of protein dynamics. Fluorescence anisotropy measurements^37–40^ and NMR relaxation^41^ have revealed that local rotational flexibility can either increase or decrease upon condensation depending on the protein and the interaction network formed within the dense phase, highlighting the heterogeneous and molecular-scale-dependent nature of condensate dynamics^56,73–75^. In Ddx4 condensates, backbone motions slow only about sixfold, whereas protein diffusion slows about 100-fold^76^. Both techniques report the reorientation of a probe or bond near a single site, a quantity different from the segment relaxation computed here. Our results could be tested directly by measuring reconfiguration times between dye pairs placed at different sequence separations, for example by nanosecond fluorescence correlation spectroscopy^21,77^, in both phases; we predict that short separations relax faster within the condensate.

This dynamical crossover cannot be attributed to local structural changes, because short segments have nearly the same size in both phases. Rather, it reflects opposite effects of the two environments. In the dilute phase, collapse slows local relaxation but not whole-chain relaxation, likely because a compact chain reorients quickly as a whole: its whole-chain relaxation time, about 40 ns, is close to the time a rigid sphere of the same size takes to reorient in water, and it follows the solvent viscosity (*Fig. 5D*). Within the condensate, neighboring chains slow whole-chain relaxation, whereas short segments relax about as fast as in an expanded chain in water (*Fig. 4F*). The dilute phase therefore sets the magnitude and extent of the speed-up: in a dense phase at fixed protein concentration, relaxation does not change with chain length or interaction strength, whereas dilute-phase relaxation does. Weakening the chain collapse in the dilute phase, by tuning protein-protein or protein-water interactions, reduces the speed-up. In addition, the crossover length grows roughly in proportion to chain length rather than matching a fixed length scale of the condensate, such as the mesh size, as it would if the speed-up arose within the condensate. Because the same net attraction drives chain collapse and phase separation^24,61^, we expect the speed-up to be largest for IDPs with a strong driving force for phase separation, at similar condensate concentrations. Furthermore, water dynamics strongly modulate the relative relaxation between dilute and dense phases, because dilute-phase relaxation is more strongly coupled to the solvent viscosity. Simulations without explicit water eliminate the speed-up entirely, causing all segments to relax more slowly within the condensate than outside it (*Fig. 6A*). This trend persists even after the solvent-free dense phase is re-equilibrated, allowing its density to readjust. Water thus plays opposite roles in the two phases: it slows the collapsed chain in the dilute phase but keeps short segments mobile within the condensate. This finding agrees with the proposal that water-mediated interactions preserve condensate fluidity^63^ and with simulations in which water loss drives condensate aging^78^.

Together, these results show that phase separation selectively accelerates short-segment protein motion, even as it slows global protein dynamics. Short segments can move quickly despite the high viscosity of the bulk condensate, which is set by its slowest, chain-scale relaxation^14,77,79–81^, as they feel only their immediate, water-rich surroundings. This coexistence of slow global dynamics with rapid local relaxation provides a possible explanation for how condensates can maintain liquid-like molecular mobility despite their large bulk viscosity.

Our results also suggest that condensates can concentrate proteins while speeding up the local segmental motions on which molecular recognition and chemical modification rely^9^. Short linear motifs, which mediate many such interactions, span only a few residues^82^, well within the range of segment lengths that relax faster in the condensate. Condensates are often thought to speed up biochemical reactions by concentrating enzymes and substrates^83,84^; faster local motions of the participating proteins may also contribute. Moreover, chain length changes whole-chain relaxation within the condensate but not local relaxation (*Fig. 2D*), so changes in protein length could tune condensate material properties without changing local protein motions. Our work emphasizes the importance of considering dynamics across multiple molecular length scales for understanding condensate material properties and biological function.

## METHODS

### Simulation Details

To study the dilute and dense phases of FUS-LC and (SYGQ)*_n_* sequences with *n* = 10, 20, 30, 40, and 50, we conducted single-chain and multichain simulations using the Martini 3 coarse-grained MD model^85^ with explicit solvent and ions. All system construction and initial preparation were performed using GROMACS^86^ (version 2022). Equilibration and production simulations were carried out in OpenMM^87^ (version 8.1) using the Martini implementation developed by MacCallum and co-workers^88^. All dilute and dense phase production simulations were carried out at 300 K in a cubic periodic simulation box with trajectories saved every 50 ps to compute conformational and dynamical properties. Nonbonded interactions were evaluated with a cutoff distance of 1.1 nm for both van der Waals and electrostatic interactions, and long-range electrostatics were treated using the reaction-field method^89^ with an effective dielectric constant of *ε_r_* = 15, the standard Martini 3 setting^85^, used for all simulations. FUS-LC and (SYGQ)*_n_* chains were coarse-grained using the martinize2 Python script^90^, starting from atomistic conformations. The simulation compositions, final box volumes, concentrations, production durations, and replica counts are summarized in *SI Table S1*.

The dilute phases were created using a single protein chain in an initial box (*L* = 18 nm), followed by solvation with Martini water. The box was then equilibrated in the *NPT* ensemble using a Monte Carlo barostat^91^. Some water beads were replaced by NaCl to reach an ion concentration of 100 mM. Production simulations were carried out in the *NVT* ensemble using a Langevin middle integrator^92^ with a timestep of 20 fs and a friction coefficient of 0.1 ps^-1^. To improve statistical sampling, 6 to 11 independent *NVT* simulations (*SI Table S1*) were initiated from distinct initial configurations. These initial configurations were extracted as snapshots from equilibrated trajectories at different time points. Each replica was simulated independently under identical conditions, and the mean and standard error of the mean (SEM) were calculated across independent replicas for all single-chain properties.

The cubic dense phases were generated by extracting the equilibrated condensate region from the slab simulations and converting it into a cubic simulation box. The resulting systems were then energy minimized and equilibrated in the *NPT* ensemble following solvation with Martini water and the addition of NaCl to achieve a target salt concentration of 100 mM. The *NPT* equilibration was performed in two consecutive stages using a Monte Carlo barostat^91^: an initial equilibration at 15 bar for 50 ns, followed by equilibration at 1 bar for an additional 50 ns. The number of peptide chains, water particles, and the final box volume were chosen (*SI Table S1*) such that all cubic dense-phase simulations maintained a protein concentration of 680 mg/mL and a corresponding water concentration of approximately 600 mg/mL consistent with the average concentrations estimated from coexistence slab simulations of FUS-LC (*SI Fig. S9* and *SI Table S1*). This concentration is based on Martini 3 bead masses, which exceed the real protein masses (24,768 vs 17,168 g/mol for FUS-LC); in real protein mass, it corresponds to about 470 mg/mL, close to the 477 mg/mL measured for FUS-LC condensates by NMR^17^. We kept this concentration fixed for all chain lengths and for the interactionstrength and water-mass variants, so that differences in dense-phase dynamics do not arise from differences in protein concentration. Following *NPT* equilibration, production simulations were carried out in the *NVT* ensemble using a Langevin middle integrator^92^ with a timestep of 20 fs and a friction coefficient of 0.1 ps^-1^. The dense-phase production trajectories were divided into two equal-length blocks, and the mean and SEM were calculated from the block averages.

### Interaction Strength and Water Mass Scaling Simulation Details

We selected the (SYGQ)_40_ dilute and dense phase systems for the interaction/water-mass rescaling study. Using the procedure proposed by Thomasen and co-workers^93,94^, we scaled the PP and PW nonbonded Lennard-Jones potential interactions^60,95–97^ in the dilute and dense phases by a factor of *λ_PP_* (= 1.00 (original force field), 0.98, and 0.96) and *λ_PW_* (= 1.00 (original force field), 1.02 and 1.05), respectively. This scaling serves only to vary interaction strength in a controlled way; Martini3-IDP, a recent alternative, instead refits the bonded parameters for disordered proteins^98^. Following the modification, we energy minimized the system and started the simulations directly in the *NVT* ensemble using the same simulation settings discussed earlier, with the dense phase kept at 680 mg/mL. The system compositions, concentrations, production durations, and replica counts for the interaction-strength variants are summarized in *SI Table S1*.

To study the effect of water motion timescale (and hence the solvent viscosity, η∼*M*^0.5^) on the faster relaxation within the condensate, we varied the mass of the Martini water in both the dilute and dense phases^64–68^. In the Martini model, a single coarse-grained water bead typically represents four water molecules and has a mass of *M* = 72 g/mol^85^ (*Fig. 5A*). We scaled this mass by a factor of *m_W_* such that the scaled mass is *M*_scaled_ = *m_W_M*. Five values of *m_W_* were tested: 1 (original force field), 2, 4, 8, and 16. Following the modification, we energy minimized the system and started the simulations in the *NVT* ensemble (at the same concentrations) using the same simulation settings discussed earlier. The corresponding system compositions, concentrations, production durations, and replica counts are provided in *SI Table S1*.

### Water-Free Simulation Details

We selected the (SYGQ)_40_ dilute and dense phase systems for water-free simulations and removed the water beads from the initial coordinates and topology files, while retaining the ions. To assess the contribution of ions to segmental relaxation dynamics, we also performed waterfree simulations in which ions were removed, leaving only the protein chains. Following these modifications, we energy minimized the systems and started the simulations in the *NVT* ensemble (at the same protein concentration of 680 mg/mL as the hydrated system). To examine the effect of retaining the original simulation volume, we additionally removed the ions from the water-free dense-phase system and re-equilibrated it in the *NPT* ensemble, allowing the box dimensions and protein density to readjust. Following re-equilibration, the protein concentration in the dense phase increased from ∼680 to ∼1596 mg/mL (*SI Table S1*). We then reintroduced Na and Cl ions based on the re-equilibrated box volume to achieve a salt concentration of approximately 100 mM, reducing the number of ion pairs from 154 in the original system to 65 in the re-equilibrated system. We analyzed the chain conformations and segmental relaxation dynamics to determine how water removal (*Fig. 6*), subsequent ion removal (*SI Fig. S7*), and density re-equilibration (*SI Fig. S8*) affected the dynamical behavior observed in the hydrated systems. Because dilute-phase dynamics are faster without water (*Fig. 6A*), we saved trajectory frames every 25 ps for both the dilute and dense phases. The remaining simulation settings were identical to those described above.

### Normal Mode Analysis

Normal modes ***X****_p_* of a chain of length *N* are defined as 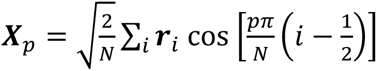 where ***r****_i_* is the position vector of the *i*-th monomer’s Martini backbone bead^14,27,47,48,99,100^. Mode number *p* corresponds to a subchain of (*N* − 1)/*p* segments. Therefore, modes 1 ≤ *p* ≤ *N* − 1 describe the internal relaxation of a chain. Specifically, *p* = 1 corresponds to the slowest chain-level dynamics. Lower normalized mode number *N*/*p* corresponds to shorter segments and higher values of *N*/*p* correspond to the larger segments. The autocorrelation function of ***X****_p_* is defined as 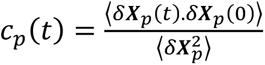 where *δ**X**_p_* = ***X****_p_* − ⟨***X****_p_* ⟩ (*Fig. 1D*; SI Figs. S1, S10, and *S11*). The *c_p_*(*t*) is described by a stretched exponential function 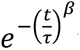 where *τ* is the characteristic timescale, and *β* is the stretching exponent. The effective relaxation time *τ_p_* of mode *p* is calculated as follows 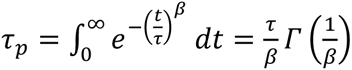 where *Γ*(*x*) is the gamma function.

## ACKNOWLEDGMENTS

This work was supported by the National Institute of General Medical Sciences of the National Institutes of Health (NIH) under grant R35GM153388, with partial support from the National Science Foundation (NSF) under grant DMR-2522693. Y.C.K. acknowledges support from the U.S. Naval Research Laboratory through the Office of Naval Research Base Program. A.N. acknowledges support by the Deutsche Forschungsgemeinschaft (DFG, German Research Foundation) through grant 470113688 and by the Klaus Tschira Foundation through grant 00.050.2024/ID 25347. We also gratefully acknowledge the Texas A&M High Performance Research Computing facility for providing the computational resources essential to carrying out the simulations reported in this study.

## DATA AVAILABILITY

All study data are included in the manuscript and/or supporting information.

## AUTHOR CONTRIBUTIONS

K.M., B.S.M., and J.M. designed research; K.M., B.S.M., D.S.D., and J.M. performed research; K.M., B.S.M., D.S.D., A.M., Y.C.K., A.N., and J.M. analyzed data; and K.M., B.S.M., D.S.D., Y.C.K., A.N., and J.M. wrote the paper.

## Supplementary Information

**Figure S1:**
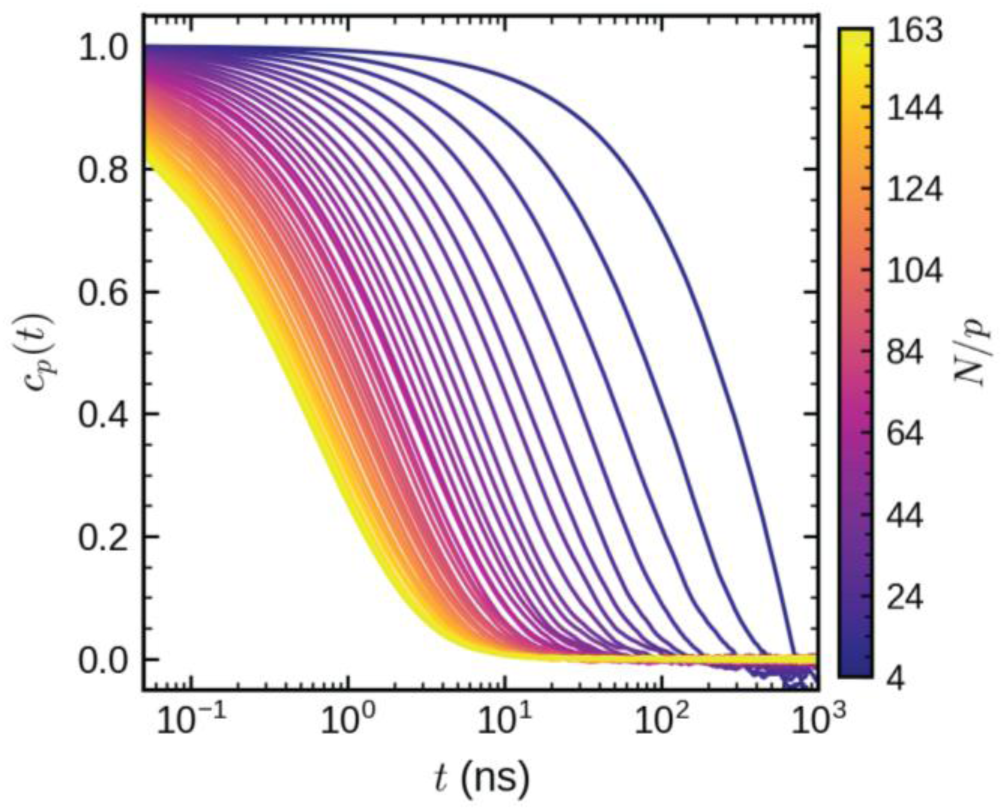
Normal modes autocorrelation function *c_p_*(*t*) for normalized mode numbers *N*/*p* = 4, 5, … , *N* of the FUS-LC sequence in the dense phase. Colors, ranging from purple to yellow, represent increasing *N*/*p*.

**Figure S2:**
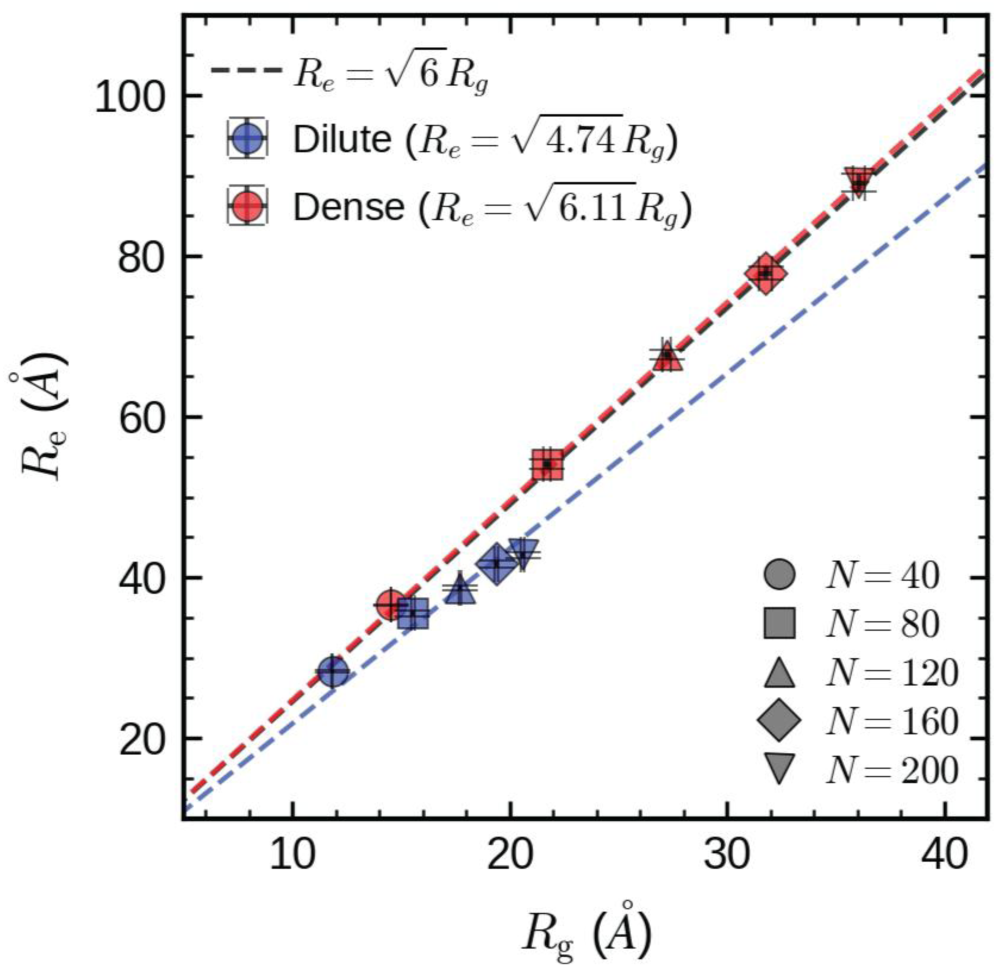
Root-mean-square end-to-end distance *R*_e_ as a function of the radius of gyration *R*_g_ for (SYGQ)*ₙ* sequences of different chain lengths (*N* = 40, 80, 120, 160, and 200) in the dilute and dense phases. Colored dashed lines represent linear fits of the form 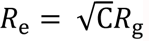, yielding C = 4.74 and 6.11 for the dilute and dense phases, respectively. The black dashed line represents the theoretical Gaussian-chain relation 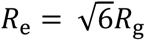.

**Figure S3:**
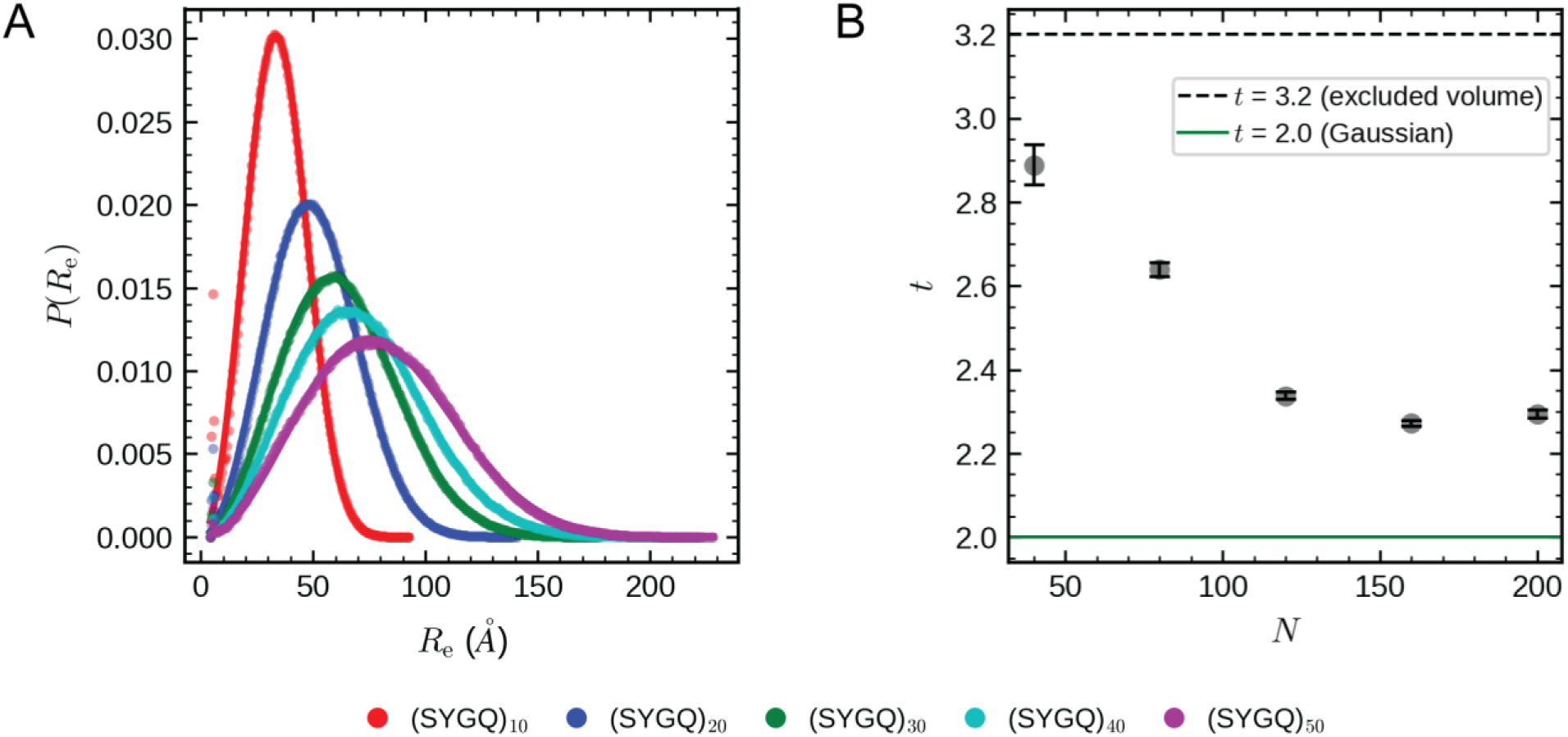
(A) End-to-end distance distributions *P*(*R*_e_) of the (SYGQ)*_n_* sequences for different chain length *N* values (*N* = 40, 80, 120, 160, and 200) in the dense phase. Solid lines represent fits to the Mazur distribution function. (B) The parameter *t* obtained by fitting the *P*(*R*_e_) of (SYGQ)ₙ sequences to the Mazur distribution function, plotted as a function of *N*. Dashed line indicates the excluded volume limit *t* = 3.2, while the solid line denotes the ideal Gaussian chain limit *t* = 2.0.

**Figure S4:**
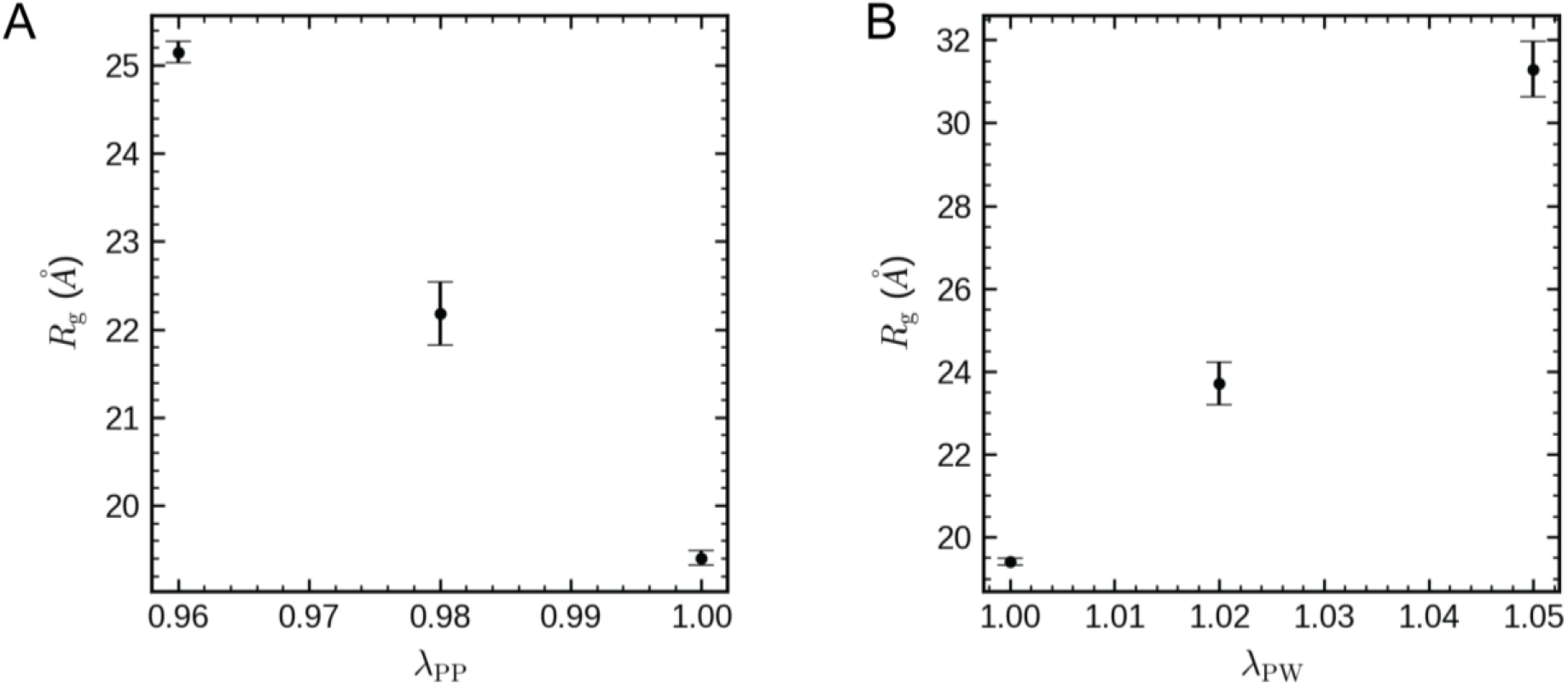
Radius of gyration *R*_g_ for the (SYGQ)_40_ sequence in the dilute phase for different (A) protein-protein (PP) interaction strengths *λ_PP_* and (B) protein-water (PW) interaction strengths *λ_PW_*.

**Figure S5:**
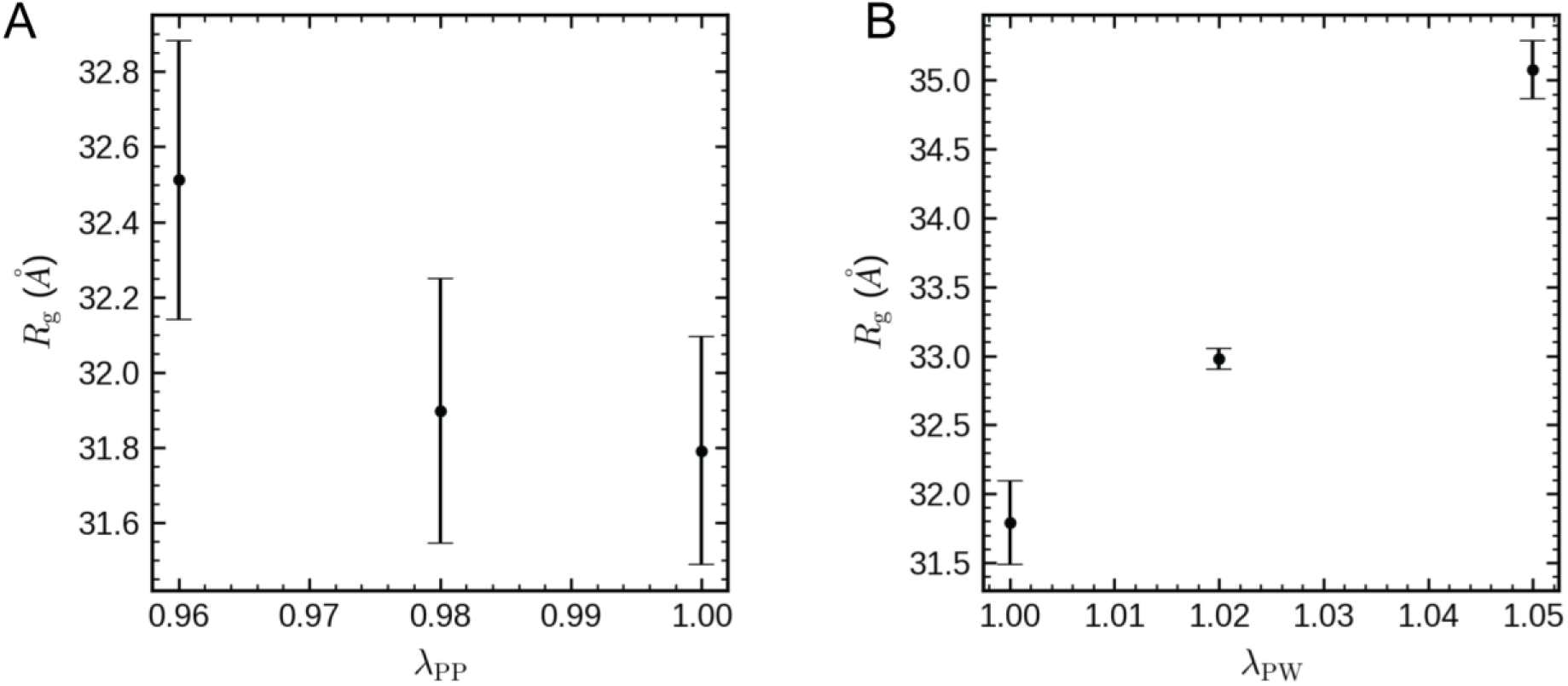
Radius of gyration *R*_g_ for the (SYGQ)_40_ sequence within the dense phase for different (A) protein-protein (PP) interaction strengths *λ_PP_* and (B) protein-water (PW) interaction strengths *λ_PW_*.

**Figure S6:**
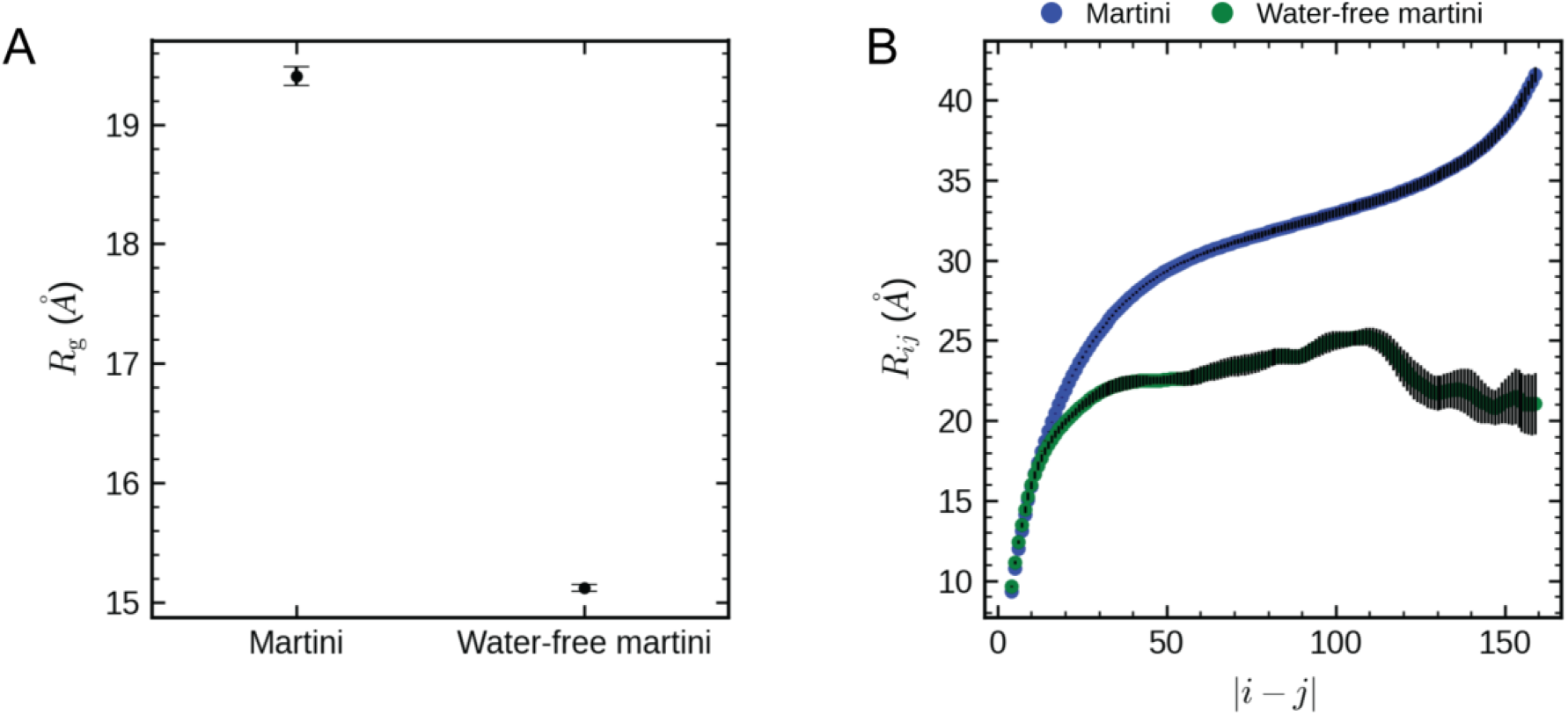
(A) Radius of gyration *R*_g_ and (B) interresidue distance *R_ij_* versus separation distances |*i* − *j*| for the (SYGQ)_40_ sequence in dilute phase for Martini and water-free Martini cases.

**Figure S7:**
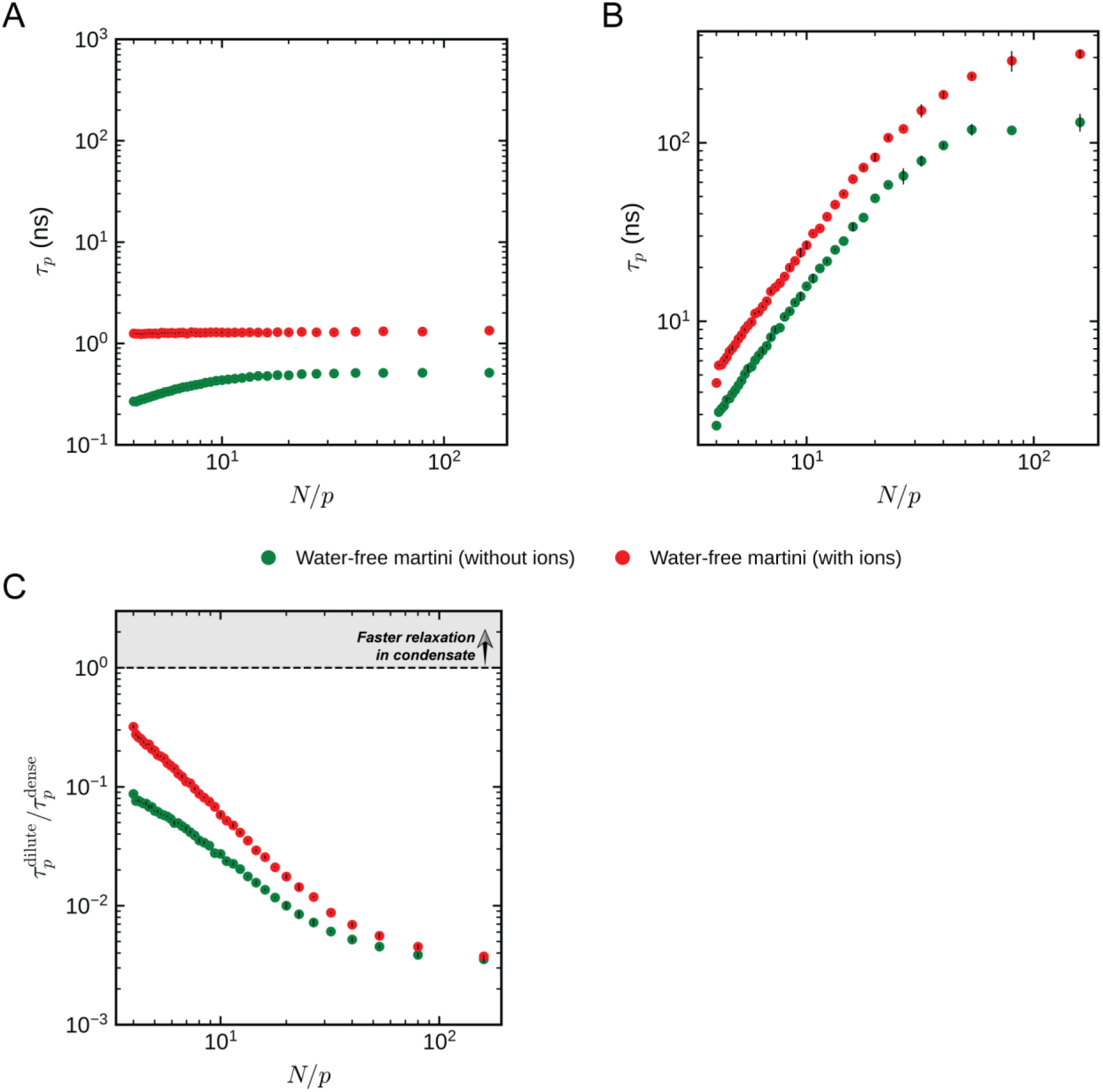
Removing Ions Does Not Restore the Speed-Up in Water-Free Simulations. Normal mode relaxation time *τ_p_* of the (SYGQ)_40_ sequence as a function of normalized mode number *N*/*p* in the (A) dilute and the (B) dense phase for Martini simulations with ions and without ions. (C) Ratio 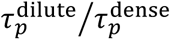 of dilute and dense phase *τp* as a function of *N*/*p* for the (SYGQ)_40_ sequence for Martini simulations with ions and without ions. The grey shaded region in (C) indicates the segments that are relaxing faster within the condensate compared to the dilute phase.

**Figure S8:**
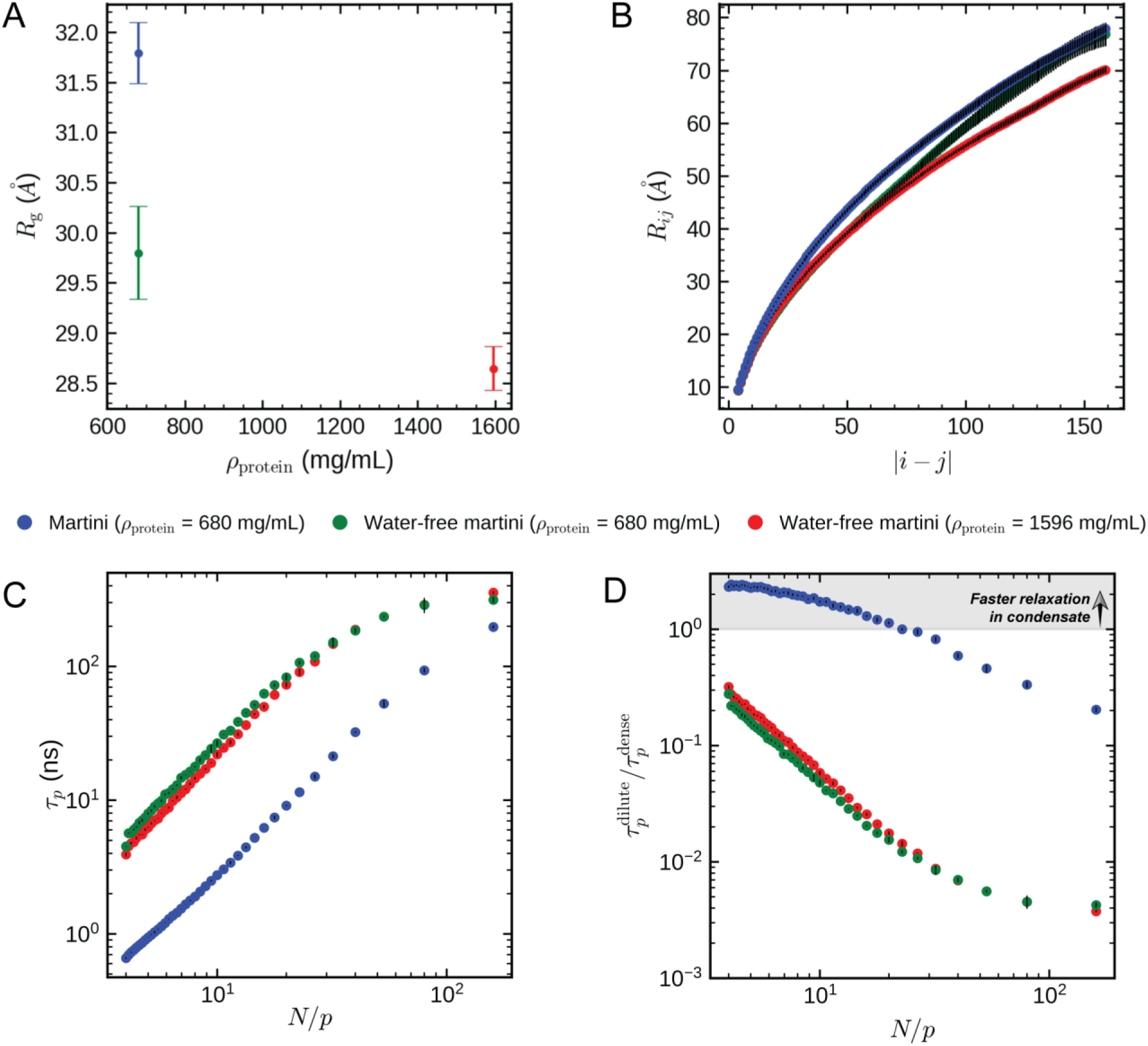
Disappearance of the Dynamical Crossover Upon Water Removal Persists Despite the Increase in Protein Density: (A) Radius of gyration *R*_g_ and (B) interresidue distance *R_ij_* versus separation distances |*i* − *j*| of the (SYGQ)_40_ sequence in dense phase for Martini and water-free Martini cases. (C) Normal mode relaxation time *τ_p_* of the (SYGQ)_40_ sequence as a function of normalized mode number *N*/*p* in the dense phase for Martini and water-free Martini cases. (D) Ratio 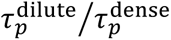 of dilute and dense phase *τp* as a function of *N*/*p* for the (SYGQ)_40_ sequence for Martini and water-free Martini cases. The grey shaded region in (D) indicates the segments that are relaxing faster within the condensate compared to the dilute phase.

**Table S1.**
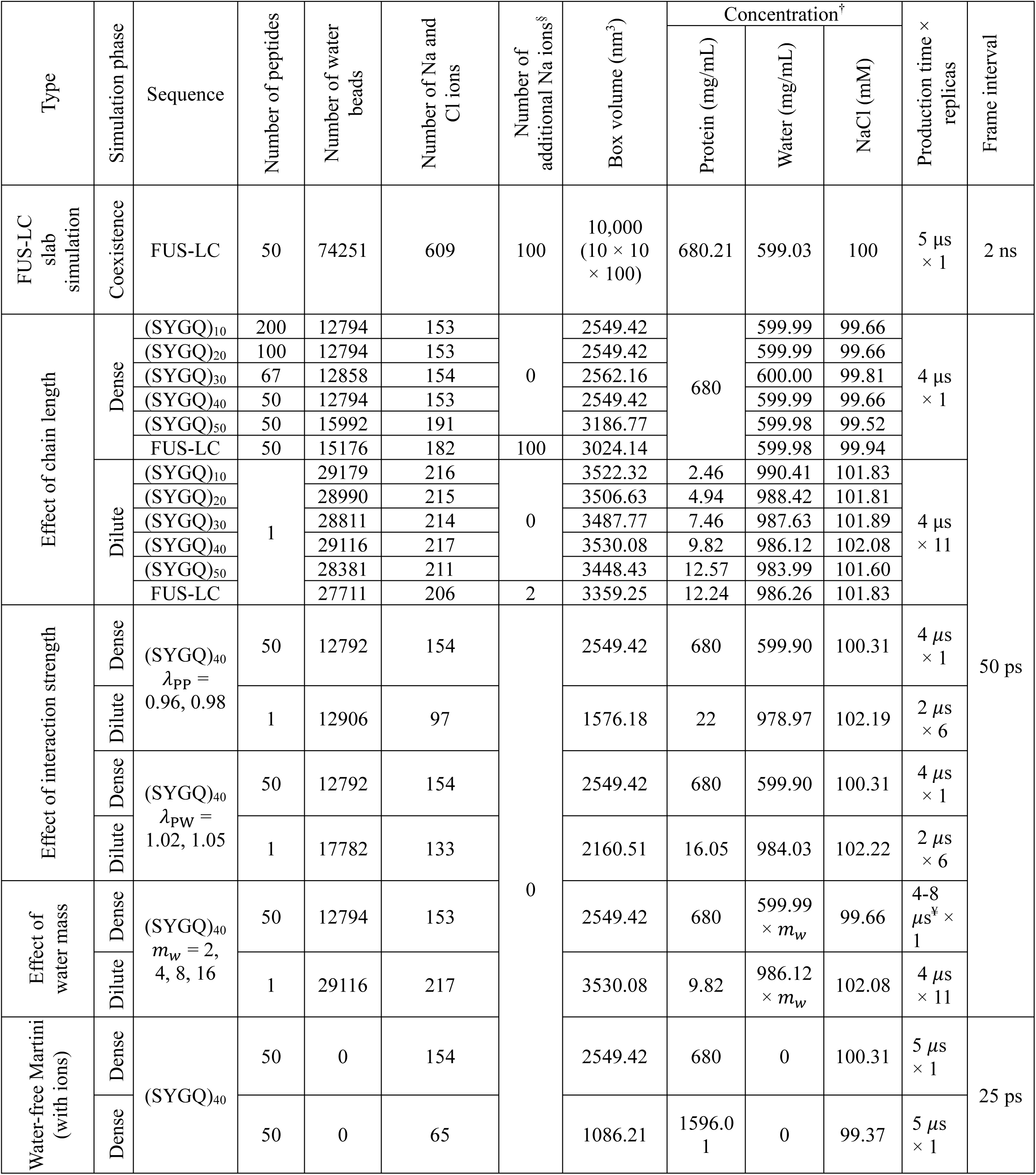
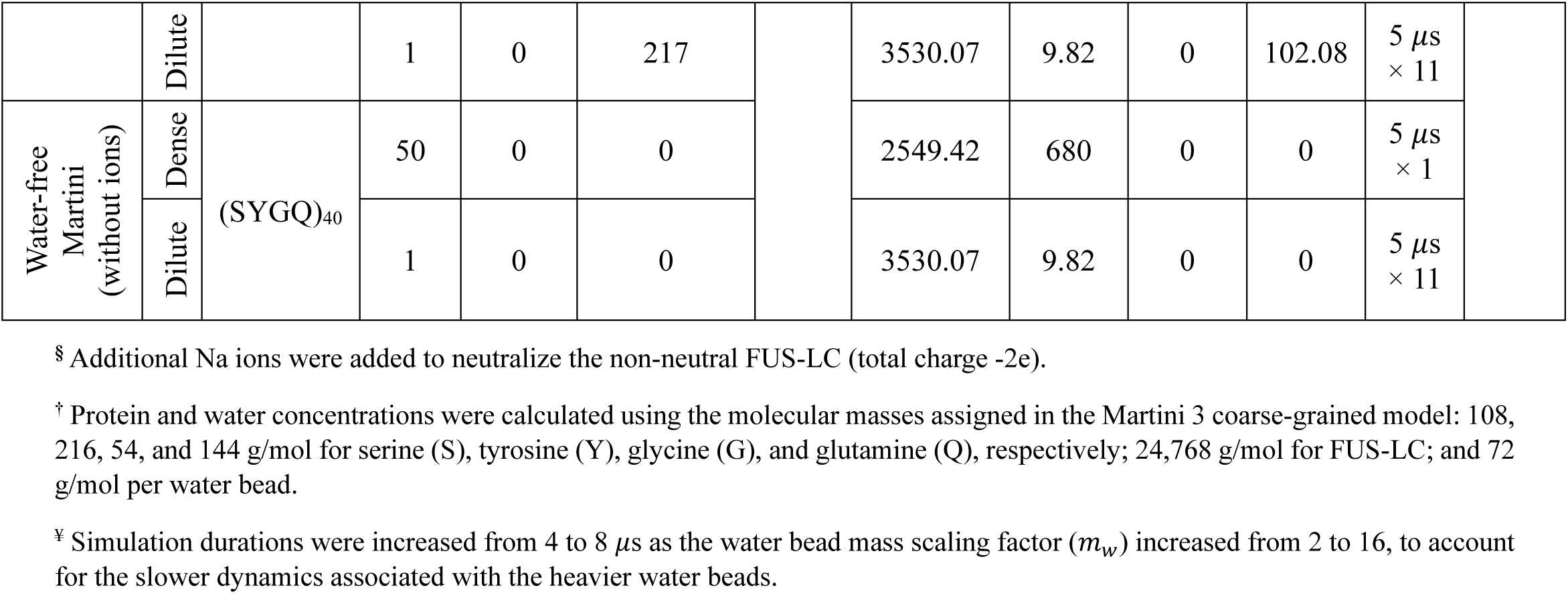
Summary of Martini 3 simulation systems and production conditions. Simulation details for the dilute- and dense-phase systems used to investigate the effects of protein chain length, protein-water and protein-protein interaction strength, and water mass scaling on protein dynamics, along with the water-free Martini systems.

| Type | Simulation phase | Sequence | Number of peptides | Number of water beads | Number of Na and Cl ions | Number of additional Na ions<sup>§</sup> | Box volume (nm<sup>3</sup>) | Concentration<sup>†</sup> |  |  | Production time × replicas | Frame interval |
|---|---|---|---|---|---|---|---|---|---|---|---|---|
|  |  |  |  |  |  |  |  | Protein (mg/mL) | Water (mg/mL) | NaCl (mM) |  |  |
| FUS-LC slab simulation | Coexistence | FUS-LC | 50 | 74251 | 609 | 100 | 10,000 (10 × 10 × 100) | 680.21 | 599.03 | 100 | 5 μs × 1 | 2 ns |
| Effect of chain length | Dense | (SYGQ)<sub>10</sub> | 200 | 12794 | 153 | 0 | 2549.42 | 680 | 599.99 | 99.66 | 4 μs × 1 | 50 ps |
|  |  | (SYGQ)<sub>20</sub> | 100 | 12794 | 153 |  | 2549.42 |  | 599.99 | 99.66 |  |  |
|  |  | (SYGQ)<sub>30</sub> | 67 | 12858 | 154 |  | 2562.16 |  | 600.00 | 99.81 |  |  |
|  |  | (SYGQ)<sub>40</sub> | 50 | 12794 | 153 |  | 2549.42 |  | 599.99 | 99.66 |  |  |
|  |  | (SYGQ)<sub>50</sub> | 50 | 15992 | 191 |  | 3186.77 |  | 599.98 | 99.52 |  |  |
|  |  | FUS-LC | 50 | 15176 | 182 | 100 | 3024.14 |  | 599.98 | 99.94 |  |  |
|  | Dilute | (SYGQ)<sub>10</sub> | 1 | 29179 | 216 | 0 | 3522.32 | 2.46 | 990.41 | 101.83 | 4 μs × 11 |  |
|  |  | (SYGQ)<sub>20</sub> |  | 28990 | 215 |  | 3506.63 | 4.94 | 988.42 | 101.81 |  |  |
|  |  | (SYGQ)<sub>30</sub> |  | 28811 | 214 |  | 3487.77 | 7.46 | 987.63 | 101.89 |  |  |
|  |  | (SYGQ)<sub>40</sub> |  | 29116 | 217 |  | 3530.08 | 9.82 | 986.12 | 102.08 |  |  |
|  |  | (SYGQ)<sub>50</sub> |  | 28381 | 211 |  | 3448.43 | 12.57 | 983.99 | 101.60 |  |  |
|  |  | FUS-LC |  | 27711 | 206 | 2 | 3359.25 | 12.24 | 986.26 | 101.83 |  |  |
| Effect of interaction strength | Dense | (SYGQ)<sub>40</sub><br>*λ*<sub>PP</sub> = 0.96, 0.98 | 50 | 12792 | 154 | 0 | 2549.42 | 680 | 599.90 | 100.31 | 4 *μ*s × 1 |  |
|  | Dilute |  | 1 | 12906 | 97 |  | 1576.18 | 22 | 978.97 | 102.19 | 2 *μ*s × 6 |  |
|  | Dense | (SYGQ)<sub>40</sub><br>*λ*<sub>PW</sub> = 1.02, 1.05 | 50 | 12792 | 154 |  | 2549.42 | 680 | 599.90 | 100.31 | 4 *μ*s × 1 |  |
|  | Dilute |  | 1 | 17782 | 133 |  | 2160.51 | 16.05 | 984.03 | 102.22 | 2 *μ*s × 6 |  |
| Effect of water mass | Dense | (SYGQ)<sub>40</sub><br>*m*<sub>*w*</sub> = 2, 4, 8, 16 | 50 | 12794 | 153 |  | 2549.42 | 680 | 599.99 × *m*<sub>*w*</sub> | 99.66 | 4-8 *μ*s<sup>¥</sup> × 1 |  |
|  | Dilute |  | 1 | 29116 | 217 |  | 3530.08 | 9.82 | 986.12 × *m*<sub>*w*</sub> | 102.08 | 4 *μ*s × 11 |  |
| Water-free Martini (with ions) | Dense | (SYGQ)<sub>40</sub> | 50 | 0 | 154 |  | 2549.42 | 680 | 0 | 100.31 | 5 *μ*s × 1 | 25 ps |
|  | Dense |  | 50 | 0 | 65 |  | 1086.21 | 1596.01 | 0 | 99.37 | 5 *μ*s × 1 |  |
|  | Dilute |  | 1 | 0 | 217 |  | 3530.07 | 9.82 | 0 | 102.08 | 5 *μ*s × 11 |  |
| Water-free Martini (without ions) | Dense | (SYGQ)<sub>40</sub> | 50 | 0 | 0 |  | 2549.42 | 680 | 0 | 0 | 5 *μ*s × 1 |  |
|  | Dilute |  | 1 | 0 | 0 |  | 3530.07 | 9.82 | 0 | 0 | 5 *μ*s × 11 |  |

<sup>§</sup> Additional Na ions were added to neutralize the non-neutral FUS-LC (total charge -2e).
<sup>†</sup> Protein and water concentrations were calculated using the molecular masses assigned in the Martini 3 coarse-grained model: 108, 216, 54, and 144 g/mol for serine (S), tyrosine (Y), glycine (G), and glutamine (Q), respectively; 24,768 g/mol for FUS-LC; and 72 g/mol per water bead.
<sup>¥</sup> Simulation durations were increased from 4 to 8 *μ*s as the water bead mass scaling factor (*m*<sub>*w*</sub>) increased from 2 to 16, to account for the slower dynamics associated with the heavier water beads.

**Figure S9.**
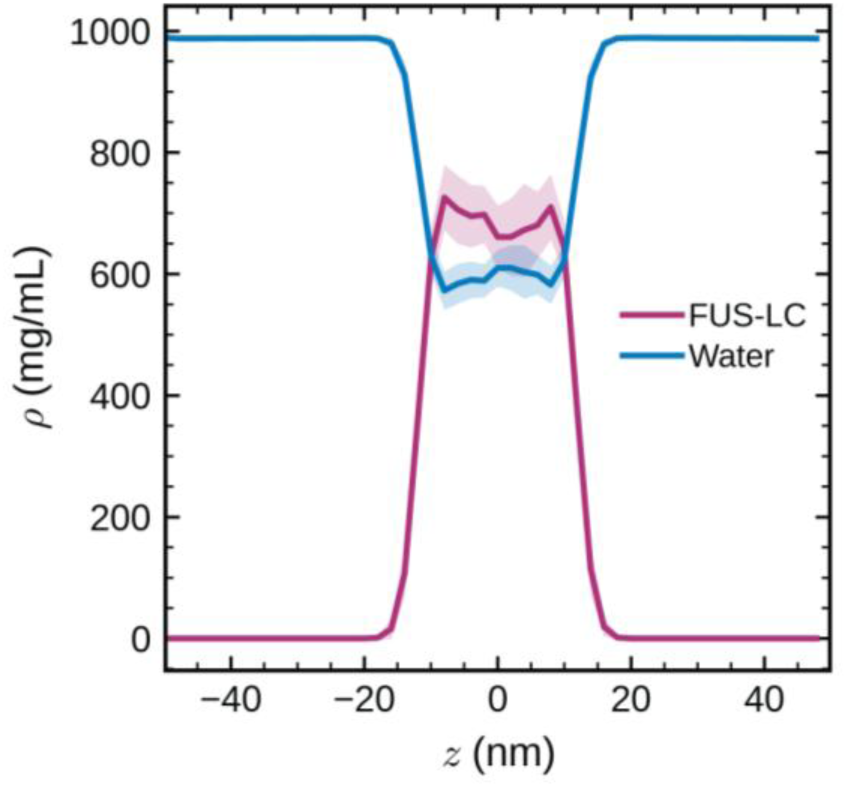
FUS-LC forms a dense slab of about 680 mg/mL protein and 600 mg/mL water. Time-averaged mass density *ρ* profiles of FUS-LC and water along the *z*-direction, obtained from slab coexistence simulation.

**Figure S10:**
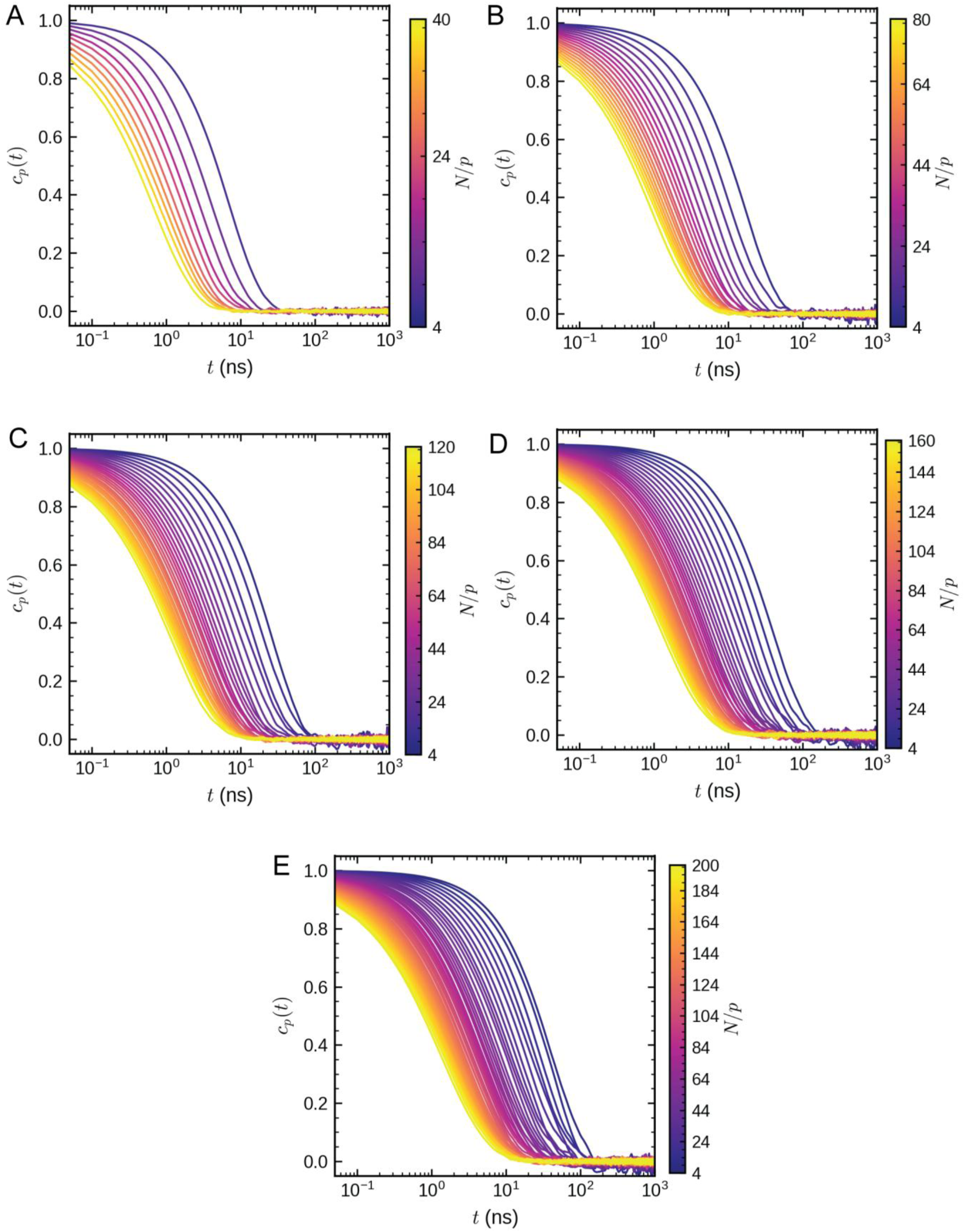
Normal modes autocorrelation function *c_p_*(*t*) for normalized mode numbers *N*/*p* = 4, 5, … , *N* of the (SYGQ)*_n_* sequences in the dilute phase for different chain length *N* values: (A) *N* = 40, (B) *N*= 80, (C) *N*= 120, (D) *N*= 160, and (E) *N*= 200. Colors, ranging from purple to yellow, represent increasing *N*/*p*.

**Figure S11:**
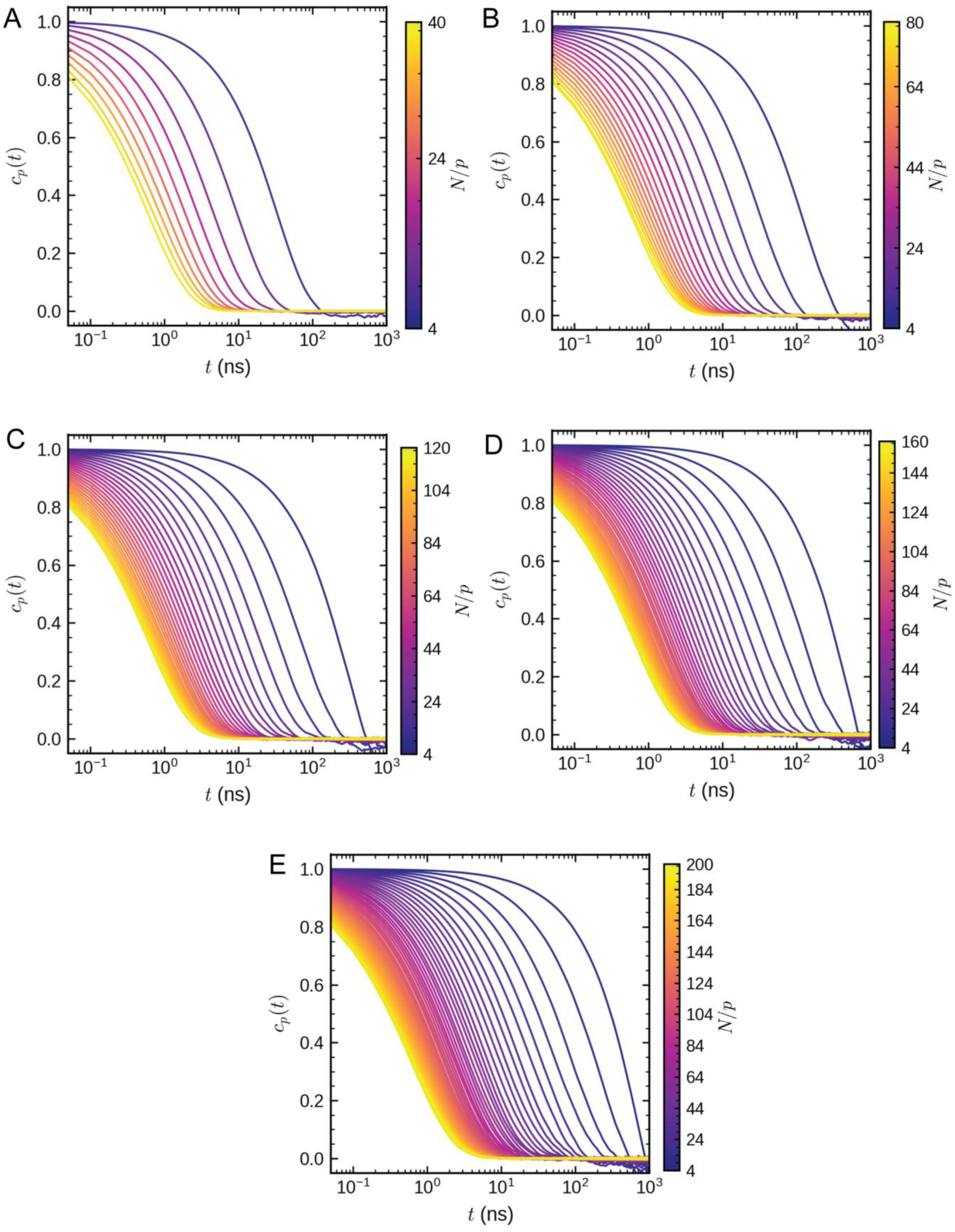
Normal modes autocorrelation function *c_p_*(*t*) for normalized mode numbers *N*/*p* = 4, 5, … , *N* of the (SYGQ)*_n_* sequences in the dense phase for different chain length *N* values: (A) *N* = 40, (B) *N*= 80, (C) *N*= 120, (D) *N*= 160, and (E) *N*= 200. Colors, ranging from purple to yellow, represent increasing *N*/*p*.

